# Continuous theta-burst stimulation of the prefrontal cortex in the macaque monkey: no behavioral evidence for within-target inhibition or neural evidence for cross-hemisphere disinhibition

**DOI:** 10.1101/2025.04.28.651079

**Authors:** Sebastian J Lehmann, Brian D Corneil

**Affiliations:** Department of Physiology & Pharmacology, Western University, London, ON, Canada, N6A 5B7; Department of Psychology, Western University, London, ON, Canada, N6A 5B7; Robarts Research Institute, Western University, London, ON, Canada, N6A 5B7

**Keywords:** TMS, continuous theta burst stimulation, pre-frontal cortex, macaque

## Abstract

**Background:** Continuous theta-burst stimulation (cTBS) can perturb neural activity and behavior by inducing effects that persist beyond the relatively short stimulation period. Although widely used in basic research and clinical settings, there lacks an understanding of the neurophysiological and behavioural effects of cTBS.

**Objectives/Hypothesis:** Two assumptions motivating the use of cTBS are that it will i) inhibit neural activity in the targeted area, and ii) consequently disinhibit neural activity in the mirroring region in the contralateral cortex. Here, we test these assumptions in the oculomotor system of healthy rhesus macaques.

**Methods:** In two macaques, we delivered cTBS between blocks of trials where they performed a delayed pro-/anti-saccade task, delivered cTBS to the right PFC (areas 8Ar and 46, which includes the frontal eye fields; 32 *cTBS-PFC* sessions), to the air as a SHAM control (27 *cTBS-SHAM* sessions), or to the nearby primary motor cortex as a brain control (21 *cTBS-M1* sessions). Across these different types of sessions, we compared changes in oculomotor behavior (reaction times, error rates, peak saccade velocity), and changes in neural activity recorded from the left, contralateral PFC.

**Results:** Despite multiple lines of evidence consistent with TMS influencing neural activity in the cTBS-PFC and cTBS-M1 sessions, we found no behavioral evidence for inhibition of the right PFC in the cTBS-PFC sessions, nor any evidence for contralateral disinhibition in the left PFC.

**Conclusions:** Our results call into question some of the fundamental assumptions underlying the application of cTBS.

**Highlights:**

- cTBS widely used in lab and clinic to rebalance activity across cortex
- We tested this in a monkey model, delivering cTBS to the prefrontal cortex
- No behavioural evidence for inhibition of brain area targeted by cTBS
- No evidence for disinhibition of spiking activity in mirroring, contralateral cortex
- Results question key assumptions about how cTBS influences network activity

## Introduction

Transcranial magnetic stimulation (TMS) is one of the most common forms of non-invasive brain stimulation. TMS is widely used in both basic research and clinical scenarios, offering a reliable means to manipulate cortical activity in a targeted and non-invasive manner. Repetitive forms of TMS such as theta burst stimulation (TBS) are particularly intriguing, as they induce neural effects that persist for some time after its application. For example, continuous theta-burst stimulation (cTBS) is thought to transiently decrease neural activity in the targeted area. Such effects were first reported in primary motor cortex (M1, Huang et al., 2005, 2017), but cTBS is now routinely applied to any number of brain areas (Franca *et al*., 2006; Rahnev *et al*., 2013; Hanlon *et al*., 2015; Valchev *et al*., 2015; Blumberger *et al*., 2018; Strzalkowski *et al*., 2019). Doing so assumes that the effects of cTBS-M1 will generalize to other brain areas, and that such effects will modulate activity within anatomically and functionally connected regions. Indeed, one common assumption motivating its use is that cTBS-induced inhibition of the targeted brain area will increase, via disinhibition, activity in the mirroring, callosal targets (Mochizuki *et al*., 2007; Stefan *et al*., 2008; Suppa *et al*., 2008, 2016; Lefaucheur *et al*., 2020). The idea that repetitive TMS can ‘rebalance’ activity across hemispheres via ipsilateral inhibition and contralateral disinhibition have motivated its exploration as a treatment mode following stroke (Zheng, Liao and Xia, 2015; Li *et al*., 2016; Long *et al*., 2018; Vink *et al*., 2023) or intractable depression (George *et al*., 2010; Li *et al*., 2014; Kang *et al*., 2016; Brunoni *et al*., 2017; Theleritis *et al*., 2017).

Despite widespread use there remains a gap in knowledge in the precise neurophysiological effects of cTBS both in the targeted area and in interconnected areas, and a series of recent results in humans have questioned the reliability of effects of repetitive TMS protocols (Hamada *et al*., 2013; Hordacre *et al*., 2017; Perellón-Alfonso *et al*., 2018; Boucher *et al*., 2021; Magnuson *et al*., 2023). Recent consensus statements (Edwards *et al*., 2024) emphasize the importance of animal models in helping bridge the gap in knowledge. We and others (Gerits *et al*., 2011; Mueller *et al*., 2014; Balan *et al*., 2017; Romero *et al*., 2019, 2022; Pouget *et al*., 2020; Klink *et al*., 2021; Lehmann and Corneil, 2022; De Lima-Pardini *et al*., 2023) have argued that the non-human primate (NHP) macaque offers an excellent animal model, given homologies in cortical microstructure, gyrification, and the anatomy and function of distributed brain networks to those found in humans. The oculomotor system can offer a particularly useful network to study in this animal model, given similarities in retinal structure and oculomotor repertoire to humans, and the extensive work in the rhesus macaque that has detailed patterns of neural activity associated with the performance of complex behavioral tasks (Lehmann and Corneil, 2022). Further, the macaque oculomotor system has been studied with many causal techniques, such as microstimulation (Robinson and Fuchs, 1969; Robinson, 1972; Bruce *et al*., 1985), cryogenic (Johnston, Lomber and Everling, 2016; Peel *et al*., 2017, 2020; Dash *et al*., 2018) or pharmacological inactivation (Sommer and Tehovnik, 1997; Dias and Segraves, 1999), or permanent lesions (Schiller, Sandell and Maunsell, 1987; Kunimatsu *et al*., 2015; Adam *et al*., 2020), providing a grounding for how non-invasive brain stimulation techniques like cTBS should influence brain and behavior.

The focus of this current study is on the prefrontal cortex (PFC), and specifically the areas rostral to and including the anterior bank of the arcuate sulcus that includes the frontal eye fields (FEF) and adjacent area 8Ar/46. This area has been extensively targeted with TMS in humans (for a review, see Vernet et al., 2014) and in macaques (Gerits *et al*., 2011; Valero-Cabre *et al*., 2012; Gu and Corneil, 2014; Mueller *et al*., 2014; Balan *et al*., 2017). These frontal areas are reciprocally connected anatomically with their callosal target (Schwartz and Goldman-Rakic, 1984), and neurophysiological results in NHPs support the notion of crossed-hemisphere inhibition (Schlag, Dassonville and Schlag-Rey, 1998; Cohen *et al*., 2010; Ramezanpour *et al*., 2024). The PFC of the rhesus macaque therefore offers an excellent platform to test the two assumptions that cTBS-PFC will i) inhibit neural activity in the targeted area, and ii) consequently disinhibit neural activity in the mirroring, contralateral PFC (Fig. 1A).

**Fig. 1.**
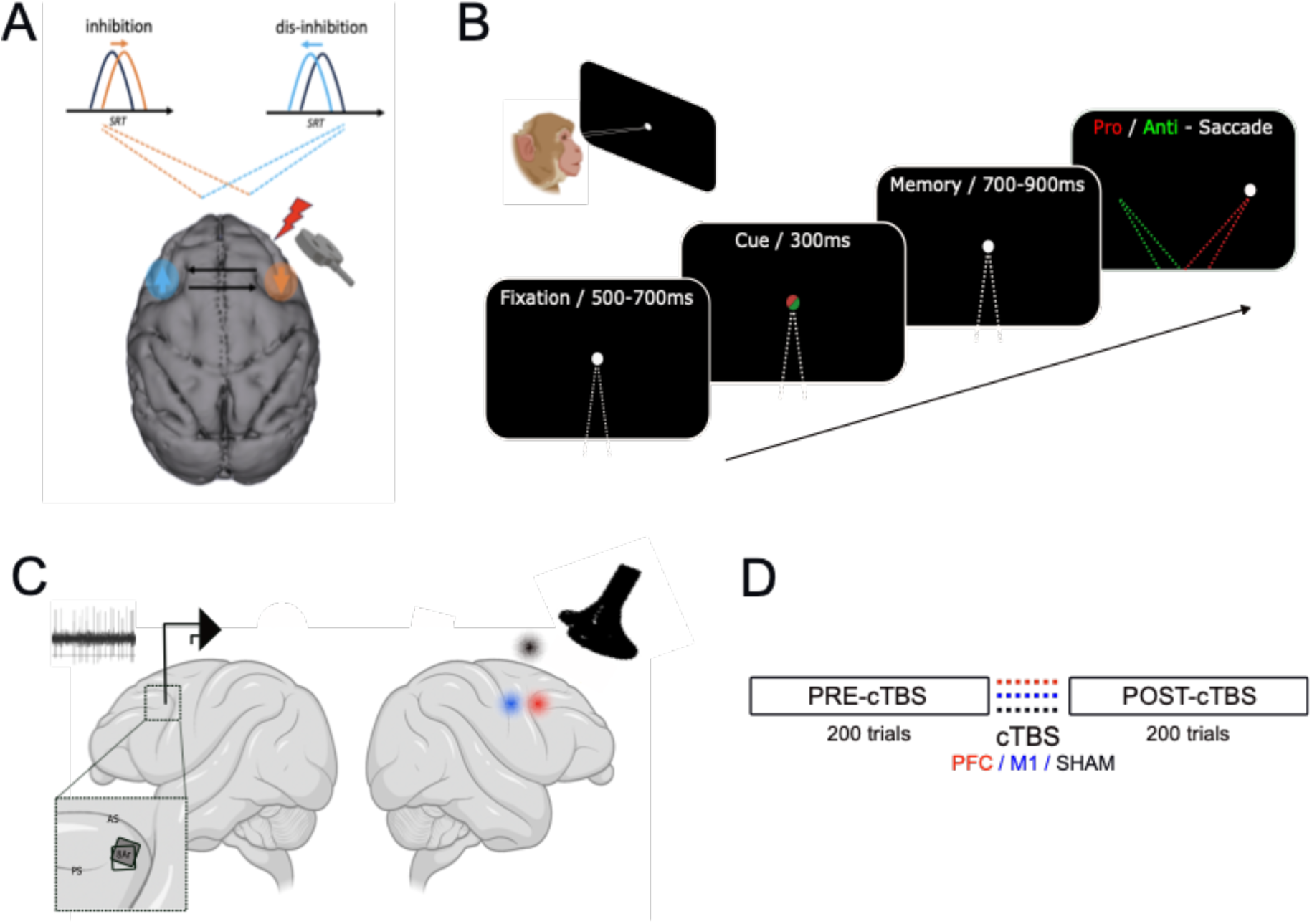
Methods. **A.** cTBS is expected to inhibit neural activity in the targeted right PFC and disinhibit activity in the contralateral left PFC, potentially impacting behavioral markers such as error rate, saccadic reaction time and peak velocity, and spiking activity in the contralateral PFC. **B.** Two rhesus macaques performed a delayed pro- and anti-saccade task with a working memory component, requiring saccades toward or away from a target appearing on the left or right, resulting in four conditions. **C.** cTBS was delivered in separate daily sessions to either the right PFC (red), right M1 (blue), or as a SHAM control (black, 2 cm above the right hemisphere). Neural activity was recorded from multielectrode arrays in the left PFC-8Ar area, rostral to the arcuate sulcus and caudal to the principal sulcus. Individual positions for monkeys Bu (opaque) and Gr (grey) are shown in the magnified section. **D.** In a blocked design, the monkeys first completed 200 trials (50 per condition) of the task (PRE-cTBS), followed by cTBS delivery, and then immediately completed a second set of 200 trials (POST-cTBS). *cTBS: continuous theta burst stimulation; SRT: saccadic reaction time; PFC: prefrontal cortex; M1: primary motor cortex; 8Ar: rhesus macaque area 8Ar; AS: arcuate sulcus; PS: principal sulcus*.

To test these predictions, we investigated the behavioral and neurophysiological effects of cTBS-PFC in two healthy rhesus macaques performing a memory-guided saccade task that required them to remember the rule to generate a pro-(look toward) or anti-(look away from) a flashed target upon disappearance of a central fixation point. Successfully executing this task requires working memory for rule maintenance, stimulus encoding, cognitive control, and visuomotor integration; neural correlates of all of these features have been reported in area 8Ar (Wallis, Anderson and Miller, 2001; Everling and DeSouza, 2005; Backen, Treue and Martinez-Trujillo, 2018; Khanna, Scott and Smith, 2020). We used a block design, comparing behavior and neural activity before and after cTBS-PFC to that observed with intervening bouts of cTBS delivered either to the air (termed cTBS-SHAM, to control for acoustic effects) or cTBS delivered to the nearby hand area of motor cortex (termed cTBS-M1, serving as a brain control that does not target the oculomotor network but controls for tactile sensations arising from TMS). The hypothesis that cTBS inhibits neural activity in the targeted PFC predicts increased error rates, and longer reaction times and reduced velocities for contralateral saccades; such effects are observed with other forms of temporary inactivation of this area (Sommer and Tehovnik, 1997; Dias and Segraves, 1999; Peel *et al*., 2014). These behavioral effects should also be greater for more cognitively-demanding anti-saccades (Pierrot-Deseilligny *et al*., 1991; Pierrot-Deseilligny *et al*., 2003; Peel *et al*., 2017). The hypothesis that cTBS-mediated inhibition disinhibits the contralateral PFC predicts increased PFC activity after cTBS in the opposite hemisphere. To test this prediction, we recorded neural activity from the PFC contralateral to the side of cTBS via a chronically-implanted array.

## Methods

### Subjects and surgical procedures

The experiments were performed in two male adult rhesus macaques (*Macaca mulatta*, animals Bu and Gr, 7 and 9 years old, weighing ca. 11 and 9 kg, respectively). Animal housing, training, surgical, and experimental procedures were approved by the Animal Use Subcommittee of the University of Western Ontario Council on Animal Care, and were conducted in accordance with the Canadian Council on Animal Care policy on the use of laboratory animals, which conforms to the guidelines laid down by the National Institutes of Health regarding the care and use of animals for experimental procedures. The NHPs’ health and weight were monitored daily. Each macaque underwent two surgeries. In the first surgery, a head post was implanted to restrain head motion during training and recording sessions, along with several fiduciary markers for neuro-navigation embedded in the acrylic implant. The implant was secured to the skull using titanium and ceramic screws, with the ceramic screws placed in the anterior part of the skull where TMS would be delivered. The thickness of the acrylic above the areas of interest for delivery of TMS was limited to ∼5 mm to ensure a minimal distance to the cortical surface (for more details see Gu and Corneil, 2014). Animals then underwent structural MRI scans to confirm cortical landmarks and locate the embedded fiduciary markers, serving for surgical planning and the use of neuro-navigation techniques for precise positioning of the TMS coil (described in detail below). In a second surgery, a multi-electrode Utah array (Blackrock Microsystems, 96 channels, electrode length 1.5mm, 10-by-10 design with 400 micrometer inter-electrode spacing) was implanted in the left (contralateral to the side of cTBS) prefrontal cortex, caudal to the posterior end of the principal sulcus and anterior to the principal sulcus (see Fig. 1C for array locations in both animals, for additional technical details see Bullock *et al*., 2017). Recordings were started after a recovery period of 2 to 3 weeks.

#### Behavioural Task

In the time between the two surgeries, the macaques were trained to comfortably sit upright in an individually adjusted primate chair, with their head restrained, facing a board of LEDs while sitting in a dark, secluded experimental room. They were trained to perform a delayed pro- and anti-saccade task, which required them to hold in working memory the rule to look either toward (*pro-saccade*) or away from (*anti-saccade*) an upcoming peripheral stimulus (Fig.1B), and then apply this rule when the peripheral stimulus was presented. The task was controlled via custom written real-time LabView programs (NI-PXI controller, National Instruments, 1kHz sampling rate), and the position of the left eye was measured by a remote eye tracking system (ETL-200, 120 Hz sampling rate, ISCAN Inc, USA). Each trial began with the presentation of a central orange LED, which the animal had to fixate for a variable period of 500-700 ms, with a tolerance of ∼3 deg radius. The color of this central LED then changed briefly for 300 ms to either red or green, which cued the monkey to plan for an upcoming pro-saccade (red instructional cue) or anti-saccade (green instructional cue). The monkey had to maintain central fixation during this time. The central LED then changed back to orange for a period of 700-900 ms, and during this time the animal had to remember the rule for the current trials while maintaining fixation on the central target. The central LED was then turned off and at the same time a peripheral red target was turned on either 15 degrees of visual angle to the left or right. The monkey was rewarded with a small amount of fluid if they executed the correct pro- or anti-saccade within 500 ms of target appearance, and remained for 200ms within the target window ∼(5 deg radius). With this experimental structure, there are four unique trial types (pro- vs anti-, stimulus presentation left or right), which were presented pseudo-randomly interleaved. Monkeys Bu and Gr performed ∼9 to 11 trials per minute, respectively. At this pace, the animals completed a block of 200 correct trials in ∼15 to 25 min, depending on their daily performance. The amount of reward was kept constant within a given experimental session.

As illustrated in Fig 1D, data on any given day were collected within an experimental session that consisted of ∼200 correctly executed trials before cTBS (∼50 of each unique trial type, termed ‘PRE-cTBS’), followed by delivery of cTBS, followed immediately by another ∼200 correctly executed trials (∼50 of each unique trial type, termed ‘POST-cTBS’). cTBS always consisted of 600 pulses of TMS, delivered in bursts of 3 pulses at 50 Hz, with an inter-burst frequency of 5 Hz (40 seconds in total). cTBS was delivered to one of three targets on any given recording day, these being: the prefrontal cortex (cTBS-PFC), the hand area of motor cortex (cTBS-M1, serving as a brain control, and a control for tactile sensations and acoustic effects), or 2 cm above the head (cTBS-SHAM, which controls for acoustic effects). In both monkeys, we initially alternated sessions of cTBS-PFC and -SHAM, introducing cTBS-M1 sessions as a second control only after collecting data from ∼10 cTBS-PFC and cTBS-SHAM sessions (e.g., the majority of cTBS-M1 data in monkeys Bu and Gr were collected ∼17 or ∼5 weeks after the start of data collection; see Fig. S1 for a summary of the timing of the recording sessions). Our rationale was to prioritize data collection from cTBS-PFC and cTBS-SHAM, in case of device failure.

#### Neuronavigation and Transcranial Magnetic Stimulation

To ensure consistent positioning of the TMS coil, we used a neuronavigation system (cortEXplore, Austria) along with structural MRI scans. For each session, the fiduciary markers embedded within the acrylic implants were used to register the monkey’s head to a 3D model obtained from the MRI scans. This allowed us to precisely track and position the TMS coil relative to the cortical target areas in real-time.

Biphasic pulses of TMS were applied via a MagStim Rapid Transcranial Magnetic Stimulator with a figure-eight coil originally designed for peripheral nerve stimulation (25 mm inner coil radius; MagStim, UK) and previously used for a variety of NHP studies (Amaya *et al*., 2010; Gerits *et al*., 2011; Valero-Cabre *et al*., 2012; Gu and Corneil, 2014; Romero *et al*., 2019, 2022). At the beginning of each session, the coil was first positioned above the hand area of the right primary motor cortex (M1) to confirm the accuracy of neuronavigation, and to confirm the repeatability of the effects of single-pulse TMS across days. Before the start of data collection, 5-10 single pulses of TMS were delivered to M1 at an output setting of 30% (monkey Bu) or 32% (monkey Gr), which reliably evoked visible thumb twitches in the contralateral (left) hand on between 80-100% of attempts (see supplementary video S1 for demonstration). After this, the TMS coil was either kept in place for cTBS-M1 sessions, or adjusted for targeting PFC (∼8mm rostral to M1, using neuronavigation), where single pulses at the same intensity were confirmed to evoke either none or less than 20% hand twitches. We estimate that the output setting used for cTBS was ∼110% relative to resting motor threshold, and the observation of qualitatively different evoked responses from the -PFC vs -M1 locations confirms a degree of focality that is consistent with previous reports (Gu and Corneil, 2014). For cTBS-SHAM control sessions, the coil was positioned above the acrylic implant, placing it on a 2cm plastic spacer which was in secure contact with the acrylic implant; in this way, TMS pulses in cTBS-SHAM sessions also delivered a mechanical sensation. In all cases, the coil was locked in place by a clamp anchored to the head post. During cTBS-delivery, the TMS coil was cooled with a combination of air suction and iced water in order to prevent overheating. To avoid potential long-term accumulation of physiological effects, cTBS-PFC or cTBS-M1 sessions were never repeated on sequential days, and we always followed a day with cTBS-PFC with either a cTBS-SHAM (usually) or cTBS-M1 (occasionally in Monkey Bu only) session, or a day where data were not collected (Supplementary Fig 1). Thus, there was a minimum of 48 hours between repeated cTBS-PFC or cTBS-M1 sessions.

#### Data acquisition and analysis

Neural activity was recorded using a 128-channel Omniplex D neural data acquisition system (Plexon Inc). Neural signals were acquired and digitized at the headstage (16 bit resolution, Plexon DigiAmp), sampled at 40 kHz per channel, with bandpass filtering applied online (300–8000 Hz). All data were stored for offline spike sorting using principal component analysis techniques (Offline Sorter, Plexon Inc), the resulting spike time information was then imported into Matlab and aligned to behavioral events. Analog data (including horizontal and vertical eye movement traces and stimulus presentation) was recorded at 1kHz resolution. Data were analyzed offline using custom-written scripts in MATLAB (MathWorks). Saccade onset (saccadic reaction time, relative to target stimulus onset) and offset were detected by applying a velocity criterion (>50 degrees per second), trials with eye blinks or unstable fixation were removed from the database. Peak saccadic velocity was defined as the maximum velocity between saccade on and offset.

#### Spiking activity

In monkey Gr, we isolated 3979 units with an average of 94.7 ± 15.2 units per session (range 63 to 118), while in monkey Bu we isolated 873 units with an average of 25.7 ± 8.2 units per session (range 12 to 40). The spiking activity of each recorded neuron was aligned to three task events on a trial-by-trial basis: rule cue onset, target onset, and saccade onset. Peristimulus time histograms (PSTHs) were computed using a causal smoothing kernel based on a gamma distribution (Baumann et al. 2009). Neurons were included for subsequent analyses if they exhibited an average firing rate of at least 1 Hz across all correct trials in at least one of those alignments. To account for variability in overall firing rates across neurons, each unit’s firing rate was normalized by dividing by the maximum average firing rate observed in any of the four task conditions (L-pro, R-pro, L-anti, R-anti). Task modulation and selectivity for each epoch were assessed using one-way ANOVAs (p < 0.01), comparing spike rates within specific analysis windows for the following condition pairs: (1) pro- vs. anti-saccades during the rule epoch (200–300 ms after cue onset), (2) left vs. right target locations during the visual response epoch (75–150 ms after target onset), and (3) leftward vs. rightward saccades during the peri-saccadic epoch (−50 to +50 ms around saccade onset).

#### Statistical analyses

Behavioral analyses of saccade parameters were conducted in Matlab (MathWorks Inc.), while statistical analyses were performed using linear mixed models (LMMs) in Jamovi (v 2.3.28). LMMs were run for behavioral and neural data combined from both monkeys to quantify main and interaction effects; the Satterthwaite method was used to estimate degrees of freedom and p-values. For significant interactions, post-hoc comparisons were made using Bonferroni-adjusted p-values to correct for multiple comparisons. LMMs assessed the effects of cTBS location (cTBS-PFC, -SHAM, -M1), trial type (pro- vs. anti-), saccade direction (left vs. right, for the visual response and saccade epochs), and monkey, with sessions (for behavioral analyses) and unit IDs (for neuronal spike rate analyses) included as random effects. In the case of the more fine-grained assessment of short-term effects on saccadic reaction time, we also assessed the factor block (pre vs post cTBS delivery).

#### Data visualization

Final figures were generated using standard matlab plotting functions and the “gramm data visualization toolbox” (Morel, 2018). Parts of the Methods figure (Fig. 1) was created using “biorender.com”.

## Results

Two rhesus macaques performed an eye movement task in which the instruction to execute a pro- or anti-saccade had to be memorized until the appearance of the visual target, which served as a “go” signal for saccade execution (Fig. 1B, see *Methods* for details). After ∼200 correct trials (i.e., ∼50 per condition), 600 pulses of cTBS were delivered to either PFC, M1 (serving as “brain-control”), or above the head (SHAM-control), followed by a second block of the same number of trials. Sessions were limited to one per day, recorded across a total of 23 (monkey Gr) and 13 weeks (monkey Bu), totalling 80 sessions where cTBS was delivered either to the right PFC (17 and 15 sessions for monkey Gr or Bu, respectively), right M1 (11/10), or to the air (a SHAM control, 16/11).

In the following, we first examine saccade error rate, reaction time, and peak velocity to assess whether there are behavioural signatures of inhibition following cTBS to the right PFC. For each of these behavioural metrics, we reasoned that bilateral saccade performance could be influenced if cTBS disrupted the encoding or maintenance of the rule to execute a pro- or anti-saccade. Alternatively, unilateral saccade performance could be influenced if cTBS to the right FEF disrupted the processing of leftward visual stimuli (L-pro or R-anti), and/or the generation of leftward saccades (L-pro or L-anti). Our hypothesis predicted that such effects would be specific for cTBS-PFC but not cTBS-M1 or cTBS-SHAM, and greater on anti- vs pro-saccade trials. For these behavioural analyses, we compare the pre-post changes observed with cTBS-PFC to those observed for cTBS-M1 and cTBS-SHAM. Following these behavioural analyses, we present an analysis of spiking activity recorded from the left PFC, segregating our analysis of neural activity in segments associated with rule maintenance, target onset, and saccade execution.

For brevity, we report only main and interaction effects involving cTBS location. We observed the expected pattern of results for the other factors (e.g., higher error rates, longer reaction times, slower peak velocities for anti- vs pro-saccades), so present these results in the Supplementary Information unless they bear on the interpretation of the effects of cTBS location.

### Behavioral analyses

#### Error rates

We derived error rates for pro- and anti-saccades in each direction, doing so separately for the pre- and post-cTBS intervals for each of the three areas targeted by cTBS (Fig. 2A&B; red and green colors depict data for pro- and anti-saccades before cTBS respectively, with darker shades for saccades to the right; grey bars show the post-cTBS intervals). As expected from the additional complexity of remembering and executing an anti-saccade, error rates in both monkeys before cTBS were ∼5-15% higher on anti- vs pro-saccade trials. The change in error rate across cTBS can be appreciated by comparing these colored bars to the grey bars immediately to the right.

**Fig. 2.**
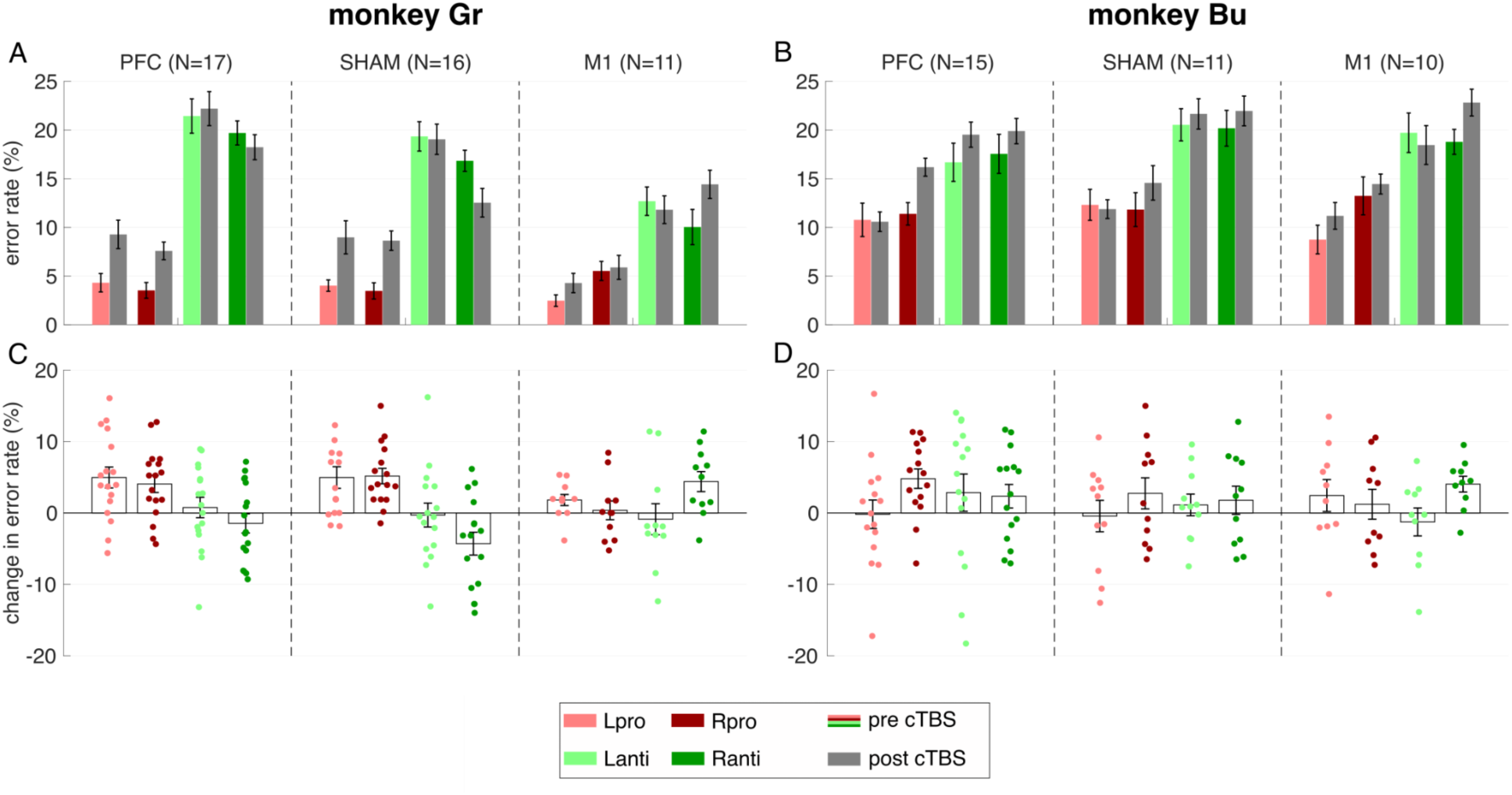
Effect of cTBS on error rates. In these and ensuing plots, the colours indicate task type (red: pro-saccades, green: anti-saccades), while lightness indicates saccade direction (light: leftwards, dark: rightwards). **A,B**. Error rates before (coloured bars) and after (grey bars) cTBS for both monkeys, averaged across sessions (error bars: SEM). **C,D**. Changes in error rate after cTBS delivery (positive values denote an increase in error rate after cTBS). Individual dots indicate the error rate change in a given session, while bar plots represent average change across sessions (error bars: SEM). *PFC: prefrontal cortex; M1: primary motor cortex; SEM: standard error of the mean*.

Our specific interest is how performance changes across cTBS, as a function of cTBS location. For every session, we therefore derived the change in error rate (Rate_post_ - Rate_pre_) across cTBS (Fig. 2C&D). Applying the linear mixed model analyses, we found that there was no main effect for cTBS location (F(2,74) = 0.52, p = 0.60; PFC = 2.26% ± 0.65, SHAM = 1.34% ± 0.72, M1 = 1.51% ± 0.80). The only significant effects involving cTBS location emerged in higher-order interactions. However, post-hoc analyses indicated that these effects were primarily driven by differences in saccade type rather than cTBS location. The significant cTBS location × direction × saccade type interaction (p = 0.004) was driven by a contrast between PFC/right/pro and SHAM/right/anti trials (p = 0.039; all other comparisons p > 0.085). Similarly, the cTBS location × saccade type × monkey interaction (p < 0.04) was driven by differences within Monkey Gr: SHAM/pro vs. SHAM/anti (p < 0.001), PFC/pro vs. PFC/anti (p < 0.042), and PFC/pro vs. SHAM/anti (p < 0.003).

Importantly, the observed changes in error rates across cTBS-PFC resembled those seen across cTBS-SHAM and cTBS-M1. Thus, there was no evidence for changes in error rate that could solely be attributed to cTBS-PFC, nor any evidence for the predicted state-dependent disruption in the animals’ ability to perform anti-saccades.

#### Saccadic Reaction Time

We present the data for saccadic reaction time (RT) in a similar manner, first showing the data recorded before and after cTBS broken down by cTBS location and monkey (Fig. 3A,B), and then by representing the change in RT across cTBS locations (Fig. 3C,D). RT data are only shown for correctly-executed trials. The expected increase in RTs for anti-saccades was more pronounced in monkey Gr (Fig. 3A) than monkey Bu (Fig. 3B). RTs tended to decrease after cTBS In monkey Bu (Fig 3B), but a similar trend was not apparent in monkey Gr (Fig. 3A).

**Fig. 3.**
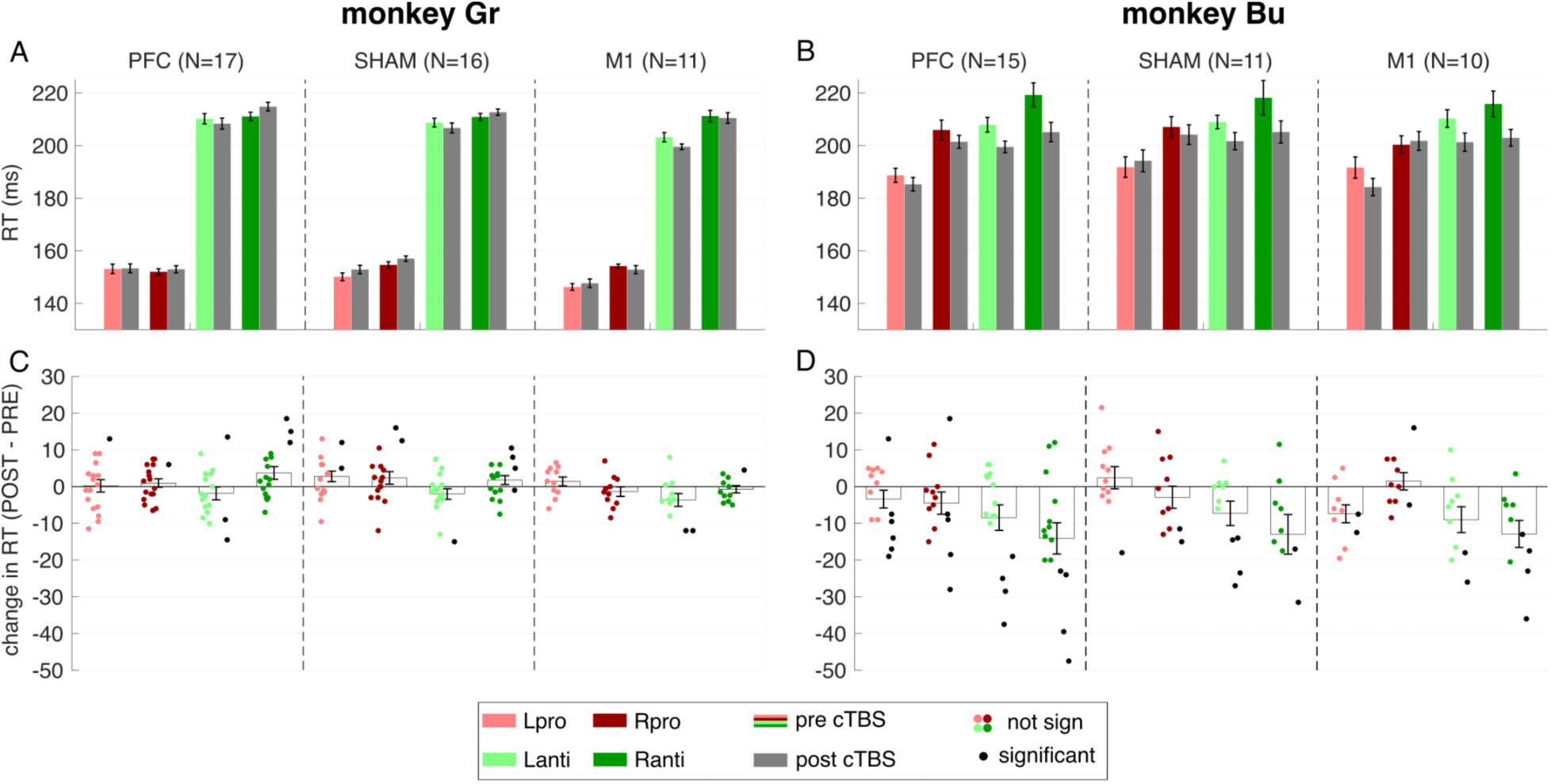
Effects of cTBS on saccadic reaction time (RT). Same general format as Fig. 2. **A,B**. RTs before and after cTBS-delivery for both monkeys, averaged across sessions. **C,D**. RT changes following cTBS delivery, relative to baseline. Black markers shifted to the right of each bar graph indicate individual sessions with significant RT changes for that condition on that day (1-way ANOVA, uncorrected), while red/green markers shifted to the left of each bar graph indicate sessions without significance. *PFC: prefrontal cortex; M1: primary motor cortex*.

As with error rate, there was no main effect of cTBS location on the change in RT across cTBS (F(2,74) = 0.52, p = 0.60; PFC = −3.43ms ± 1.27; SHAM = −1.98ms ± 1.40; M1 = - 4.02ms ± 1.56). There were also no significant two-way interactions involving cTBS location (all p > 0.05). Furthermore, post-hoc analysis of a significant interaction effect for cTBS location × direction × monkey (p = 0.03) revealed it was not driven by cTBS location but by a significant difference between monkeys (PFC/right/monkeyGr vs PFC/right/monkeyBu, difference = −11.64ms, SE = 2.79, t(108) = −4.17, p = 0.004, Bonferroni-corrected). Thus, we observed little no evidence for our prediction that cTBS-PFC should selectively lead to increased RTs, finding in fact that RTs in monkey Bu consistently decreased after cTBS regardless of stimulation location.

#### Sessions with significant RT changes across cTBS

Recent studies suggest that the effects of cTBS can vary across days within the same individual, likely due to state-dependent factors (Boucher *et al*., 2021; Ozdemir *et al*., 2021). We therefore assessed whether the proportion of sessions with significant within-session RT changes depended on cTBS location. Significant RT changes are indicated by black markers lying above or below the x-axis in Fig. 3C–D, representing sessions with increased or decreased averaged RT post-cTBS, respectively (see Table 1 for details).

**Table 1.** Proportion of sessions showing significant or non-significant changes in saccadic reaction time (RT) following cTBS, along with corresponding rounded percentages of significant sessions. *PFC: prefrontal cortex; M1: primary motor cortex; sign: significant; ns: not significant*

|  | **monkey Gr (Fig 3C)**<br>*No of sessions sign / ns*<br>*( % sign fractions)* |  |  |  |  |  | **monkey Bu (Fig 3D)**<br>*sign / ns*<br>*( % sign fractions)* |  |  |  |  |
| --- | --- | --- | --- | --- | --- | --- | --- | --- | --- | --- | --- |
|  | *Lpro* | *Rpro* | *Lanti* | *Ranti* | *total* |  | *Lpro* | *Rpro* | *Lanti* | *Ranti* | *total* |
| **PFC** | 1 / 16<br>(6%) | 1 / 16<br>(6%) | 3 / 14<br>(18%) | 3 / 14<br>(18%) | **8 / 60**<br>**(12%)** |  | 6 / 9<br>(40%) | 5 / 10<br>(33%) | 4 / 11<br>(27%) | 4 / 11<br>(27%) | **19 / 41**<br>**32%** |
| **SHAM** | 2 / 14<br>(13%) | 2 / 14<br>(13%) | 1 / 15<br>(6%) | 4 / 12<br>(25%) | **9 / 55**<br>**(14%)** |  | 1 / 10<br>(9%) | 2 / 9<br>(18%) | 4 / 7<br>(36%) | 3 / 8<br>(27%) | **10 / 34**<br>**23%** |
| **M1** | 0 / 11<br>(0%) | 0 / 11<br>(0%) | 2 / 9<br>(18%) | 1 / 10<br>(9%) | **3 / 41**<br>**(7%)** |  | 2 / 8<br>(20%) | 2 / 8<br>(20%) | 2 / 8<br>(20%) | 4 / 6<br>(40%) | **10 / 30**<br>**25%** |

In monkey Gr, significant RT changes were rare (12%, 14%, and 7% of sessions for cTBS-PFC, SHAM, and M1, respectively), with a trend toward more effects in anti-saccades, but no association with cTBS location (Chi-Square test, χ²(2, N = 176) = 1.38, p = 0.50). In contrast, monkey Bu showed more frequent RT changes (32%, 23%, and 25% for cTBS-PFC, SHAM, and M1), although though the slightly higher proportion following PFC stimulation did not reach significance (χ²(2, N = 144) = 1.15, p = 0.56).

#### Exploring cTBS-induced short-term changes in saccadic RT

The behavioural effects of cTBS may last for only a few minutes (Huang *et al*., 2005). Figure 4A&B (top rows) shows the trends of single-trial RTs relative to the time of cTBS delivery (“trial zero”), covering the 200 trials within the PRE- and POST-sessions (recall that trial types were pseudo-randomly interleaved, thus there were 50 trials of each type before and after cTBS; the colored lines represent data from the three different cTBS locations). For both monkeys, RTs tended to be shorter immediately after cTBS delivery, regardless of cTBS location. These changes were smaller for monkey Gr (ranging around −5ms across conditions, Fig 4A, top row) than monkey Bu (ranging around −20ms, Fig 4B, top row). Further, RTs tended to be more stable through time in monkey Gr, whereas RTs in monkey Bu tended to increase through both the PRE-cTBS and POST-cTBS intervals. Such increasing RTs through time are apparent for all cTBS locations, and all trial types. To analyze these short-term effects, we examined the distribution of RTs immediately before or after cTBS, only including 25 trials of each condition (opaque grey boxes in the upper row of subplots in Fig. 4), which span ∼10 minutes of data.

**Fig. 4.**
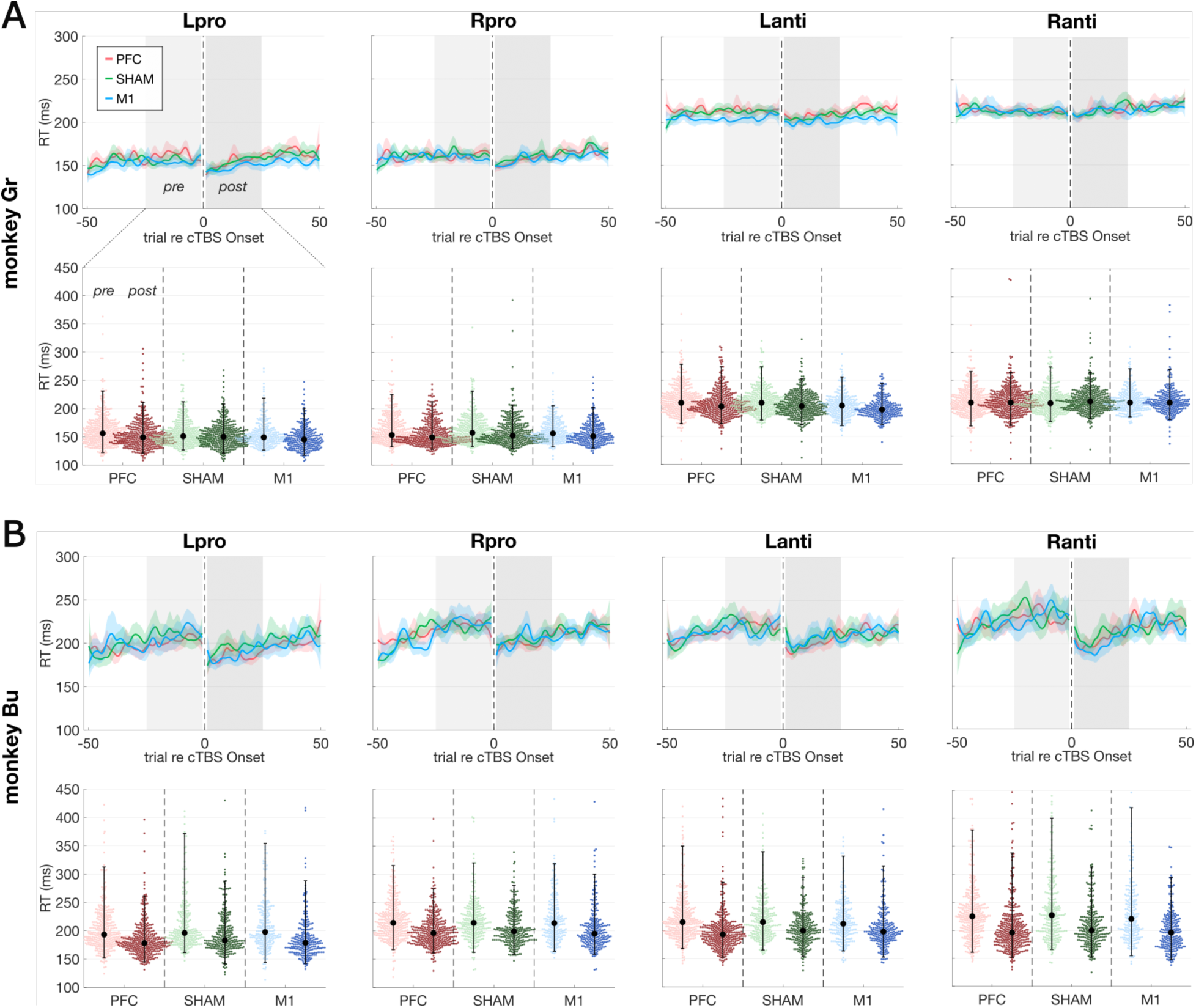
Short-term effects of cTBS on reaction time. Top row (A,B): RT trends over time relative to cTBS delivery (trial 0), shown as mean ± 95% CI across all sessions and cTBS locations (colors), for monkey Gr (A) and monkey Bu (B). Shaded grey areas demarcate the 25 trials before (light) and after (dark grey) cTBS. Bottom row (A,B): Colored dots represent individual RTs within the pre- and post-cTBS windows, with light and dark shades indicating trials before and after cTBS, respectively. Large black dots and error bars show the median and 95th percentile range. *PFC: prefrontal cortex; M1: primary motor cortex; RT: reaction time; ms: milliseconds*.

The bottom rows of Fig. 4A&B depict individual RTs from the +/− 25 trials before (light colors) or after (dark colors) cTBS delivery, across all conditions. To analyze short-term effect of cTBS location on saccadic RT, our linear mixed model included the factor block (pre- and post-cTBS), in addition to the factors cTBS-location, trial direction, trial type, and monkey.

Saccadic reaction times tended to be shorter in the post-cTBS vs pre-cTBS blocks (factor block, F(1,15921) = 558.2, p < 0.001; PFC/pre/cTBS 204ms +/− 1.3, PFC/post 190ms +/− 1.3, SHAM/pre 204ms +/− 1.4, SHAM/post 192ms +/− 1.3, M1/pre 202ms +/− 1.4, M1/post 188ms +/− 1.4), regardless of cTBS location or saccade condition. We found a significant main effect for cTBS location on RT in general (F(2,8063) = 6.98, p <0.001; PFC = 197ms ± 1.2; SHAM = 198ms ± 1.3; M1 = 195ms ± 1.3). However, post-hoc tests revealed significant differences between SHAM and M1 conditions (difference 2.7ms +/−0.7, p<0.001 Bonferroni corrected), while the differences between PFC and M1 (diff 1.5 ms, p=0.14) as well as PFC and SHAM (diff 1.22ms, p=0.23) did not reach significance. Most importantly, since our primary focus was on site-specific effects on RT following cTBS delivery, we observed no interaction effect between cTBS location and block (F(2,15921) =1.9, p=0.15), indicating that the change in RT following cTBS was not influenced by cTBS location. Post-hoc comparisons following the significant cTBS location × direction × monkey interaction (F(2,15921) = 8.96, p<0.001) revealed that most significant effects were due to differences between saccade directions and monkeys. However, we identified three significant effects related to cTBS location: PFC/left/monkeyGr vs M1/left/monkeyGr, p<0.01; SHAM/left/monkeyGr vs M1/left/monkeyGr, p=0.001; PFC/left/monkeyBu vs SHAM/left/monkeyBu, p=0.023. Similarly, post-hoc comparisons of the significant cTBS location × direction × type × monkey interaction (F(2,15921) = 3.17, p=0.042) revealed that the effect was largely driven by task-related features - specifically, strong main effects and lower-order interactions involving the factors direction and monkey. These interaction effects suggest that cTBS location contributed to the observed differences under specific conditions, but was not the primary driver of the interactions. Notably, neither the significant lower-nor higher-order interaction effects involved the factor block, which was the primary factor of interest based on our hypotheses.

In summary, we conducted a thorough analysis of the change in RTs across cTBS location, analyzing average changes in RT, the proportion of significant within-session changes in RT, and the time course of RTs. We did not find any evidence that changes in RT following cTBS delivery could be specifically attributed to cTBS-PFC, and did not observe any evidence for the predicted increase in RT that should have resulted from a hypothesized cTBS-mediated inhibition of the PFC.

#### Saccadic peak velocity

We conducted similar analyses of peak saccade velocity, and did not find any evidence for a main or interaction effects of cTBS-location (all p>0.09), nor any consistent trend for a decrease in saccade velocity predicted by cTBS-mediated inhibition of the right PFC. A figure (Supplementary Fig. S2) and further statistical analyses for task related effects on saccade velocity are included in the Supplementary Information.

### Analyses of neural spiking activity

#### Description of functional tuning before cTBS

Despite the absence of behavioral effects that conform to our predictions, cTBS-PFC may influence neural activity in more subtle ways, perhaps below the threshold required for a behavioral change or by inducing both excitatory and inhibitory effects. To explore this, we analyzed neural activity from Utah arrays chronically implanted in the frontal cortex contralateral to the cTBS site. Our analysis assesses changes in neural activity induced by cTBS within a session, comparing activity relative to the baseline preceding cTBS. Given the stability of unit recordings over time with Utah arrays (Sponheim *et al*., 2021), many units were likely recorded across multiple sessions.

A large proportion of units exhibited task modulation, and we focused on neural activity during: (1) the “rule epoch” following rule instruction, (2) the “visual response epoch” shortly after target onset, and (3) the “peri-saccade epoch” around saccade execution. Figure 5A-C illustrates three example neurons exhibiting tuned activity in these three time periods before cTBS, with the shaded areas illustrating the analysis interval. The neuron in Figure 5A exhibited higher activity for anti-saccades (green) compared to pro-saccades (red), which persisted even after removal of the instruction cue, consistent with previous reports of rule maintenance in PFC (Wallis, Anderson and Miller, 2001; Everling and DeSouza, 2005). The neuron in Figure 5B responded more strongly to visual targets in the contralateral field, with essentially identical responses on R-pro and L-anti trials. The neuron in Figure 5C exhibited higher activity before rightward (contralateral) saccades, with similar increases for R-pro and R-anti trials.

**Fig. 5.**
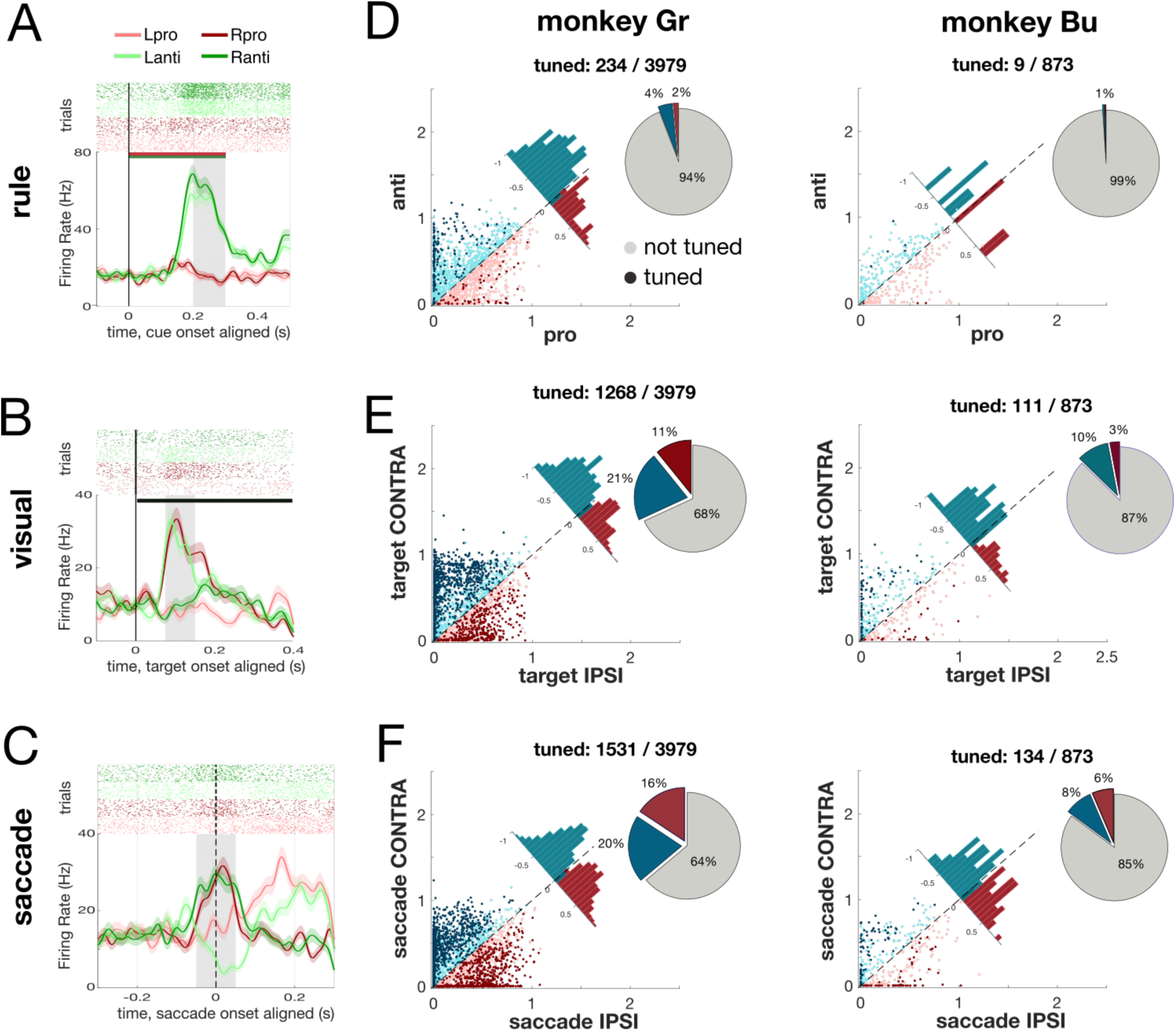
Task specific activity: single unit examples and population tuning. **A–C** Three examples of task-modulated neurons recorded in area 8Ar, showing a unit preferentially active for anti-saccades (A, aligned to cue onset, horizontal bars depict time of rule instructions), a unit with a visual response to contralateral targets (B, R-pro & L-anti, aligned to target onset), and a unit exhibiting greater activity before contralateral saccades (C, R-pro & R-anti, aligned to saccade onset). Shaded grey areas denote the analysis epochs. **D–F** Population tuning across both monkeys (monkey Gr, left; monkey Bu, right), showing the distribution of tuning directions and the percentage of neurons significantly modulated during each epoch (D: rule epoch, E: visual response epoch, F: saccade epoch). Points represent normalized firing rates of individual neurons (in arbitrary units), with darker shades indicating a significant modulation (1-way-ANOVA, p<0.01), and colors denoting tuning direction. Histograms display the distribution of tuning directions for significantly modulated units, while pie charts indicate the proportion of tuned units relative to the total recorded population in each monkey.

Figure 5D-F shows population tuning properties for all neurons recorded in the two monkeys, based on their activity in these epochs (dark points indicate neurons where activity was significantly modulated during the respective epoch using a 1-way ANOVA, p<0.01). Of the 3979 neurons recorded in monkey Gr (left column in Fig. 5D-F), 6% of neurons exhibited rule-tuning (4% for anti-saccades), 32% exhibited visual tuning (21% for contralateral targets), and 36% were tuned for saccade direction (20% for contralateral saccades). Out of the 873 neurons recorded in monkey Bu (Fig. 5D-F, right), only 9 units (1%) were rule-tuned, 13% exhibited visual tuning (10% for contralateral targets), and 15% were tuned for saccade direction (8% for contralateral saccades). These preferences for contralateral tuning to visual targets and more balanced contra- vs- ipsilateral tuning to saccade direction resembles previous findings in area 8Ar (Bullock et al., 2017).

#### Assessment of changes in task-related activity over time

Neural activity in the PFC can vary during a session due to fluctuations in slow-varying factors like attention, arousal, and motivation (Cowley *et al*., 2020). Given our blocked design, such longer-timescale variations may mask any changes induced by cTBS. Thus, we first examined changes in task-related activity across the entire session, and used this to define the interval over which we analyzed changes induced by cTBS. Figure 6 shows the average activity of significantly task-modulated units in each epoch, aligned to the time of cTBS delivery (trial 0). In both monkeys, we generally observed a gradual decrease in neural activity, particularly in the visual (Fig. 6B, E) and saccade (Fig 6C, F) epochs; the exception is the general increase in the activity of neurons encoding the anti-saccade rule in monkey Gr (Fig. 6A, right panel). Over the timeframe of the 200 trials before and after cTBS, these slow fluctuations could change normalized neural activity by ∼10-20%, regardless of cTBS location. To reduce the influence of these non-cTBS-specific, time-dependent changes while preserving sufficient trial numbers, we focused subsequent statistical analyses on the 25 trials immediately before and after cTBS (highlighted in grey in Fig. 6; these intervals span ∼10 minutes each). We also conducted the same analysis across the full set of trials (not shown), but this did not provide any additional insights beyond what is reported below.

**Fig. 6.**
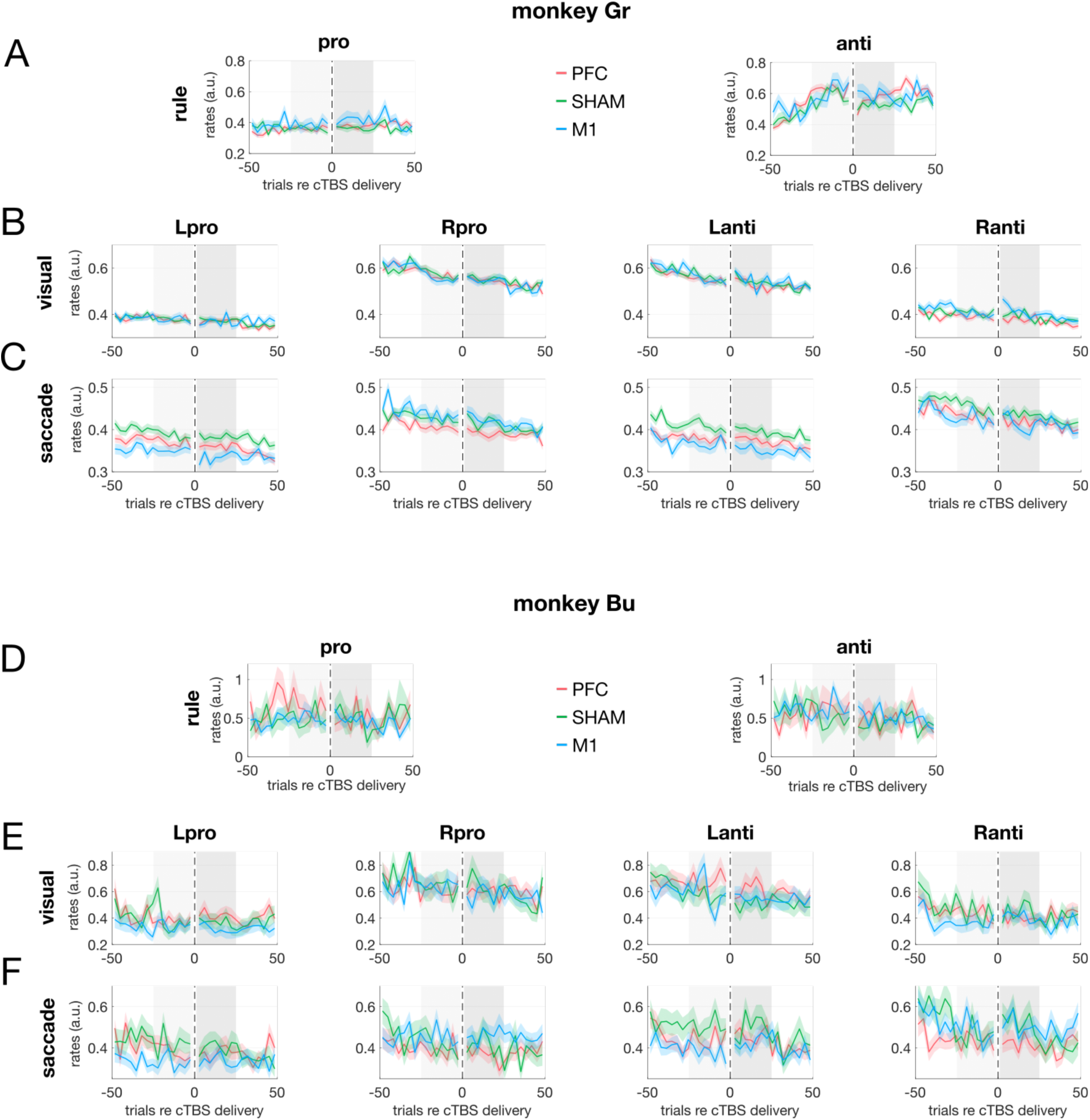
Normalized spiking activity of task modulated neurons over time. Average normalized spike rates for all significantly task-modulated units, aligned to the time of cTBS delivery (trial 0), shown separately for monkey Gr (A–C) and monkey Bu (D–F). Each row corresponds to a different task epoch: rule (A, D), visual (B, E), and saccade (C, F). For the rule epoch, neural activity is grouped by saccade type (pro vs. anti); for the visual and saccade epochs, all four task conditions are displayed. Line colors indicate cTBS location (PFC, SHAM, M1), shaded colored areas represent SEM. Grey shaded areas indicate the time windows used for subsequent statistical analyses of cTBS-induced effects. Y-axis scales (in arbitrary units) vary between epochs and monkeys to best illustrate condition-specific trends. *PFC: prefrontal cortex; M1: primary motor cortex; SEM: standard error of the mean; a.u.: arbitrary units*.

#### Population analysis of cTBS effects on spike rates

Figure 7 shows the change in average spike rate across cTBS for every neuron that exhibited significant tuning during the various epochs of the task. Given that cTBS was delivered to the other hemisphere, contralateral disinhibition predicted increased neural activity following cTBS-PFC, i.e., the red distributions in Fig. 7 should shift upward. For the rule epoch, the linear mixed model included the factors cTBS location and trial type (pro- vs. anti-saccades, i.e., the only available information at that epoch), while analyses for the visual and saccade responses also include saccade direction.

**Fig. 7.**
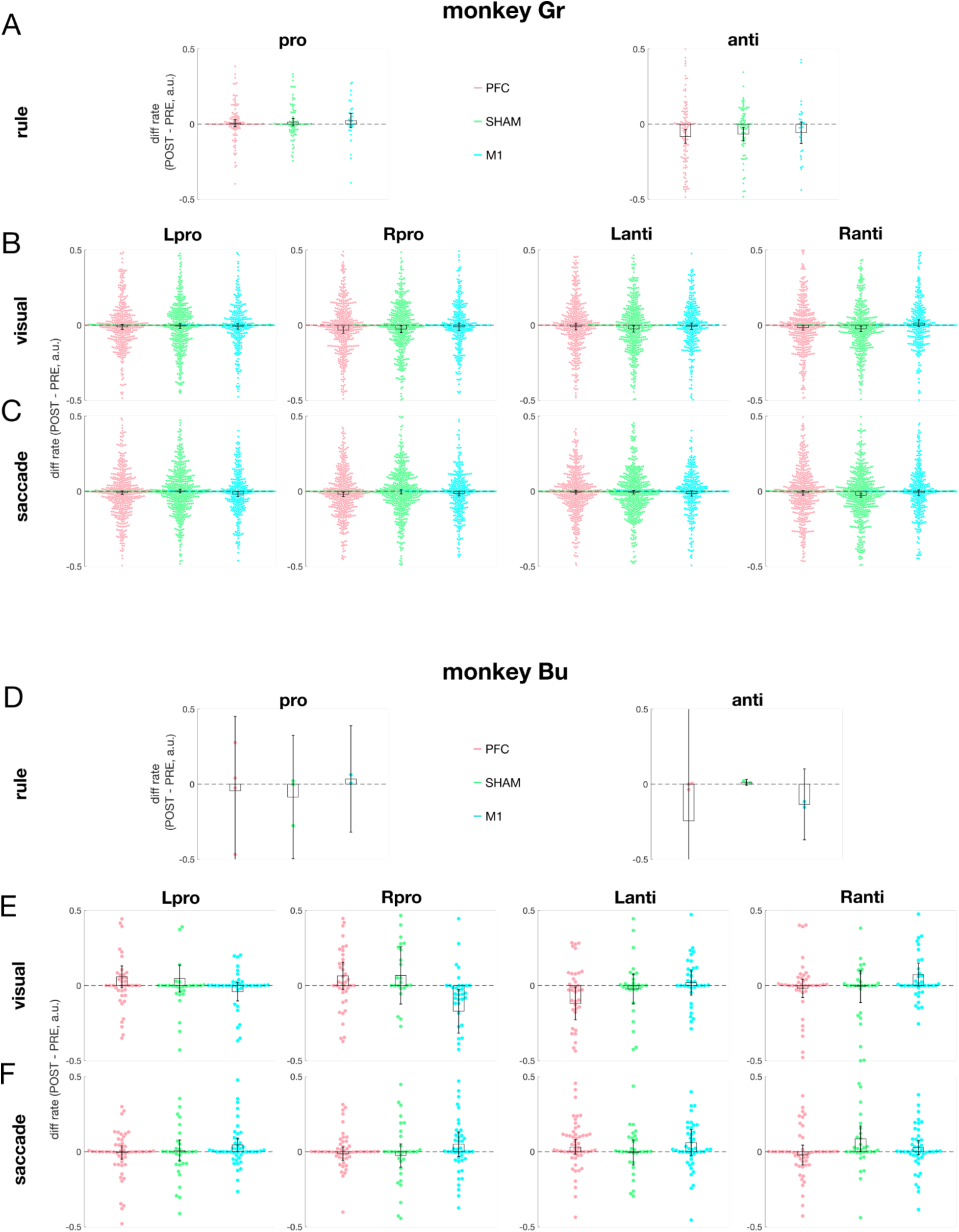
Effects of cTBS on normalized spike rates of individual neurons. Differences in spiking (post– pre cTBS, in arbitrary units) for all significantly task-modulated neurons in each epoch, shown for monkey Gr (A–C) and monkey Bu (D–F). Rows correspond to task epochs: rule (A, D), visual (B, E), and saccade (C, F). For the rule epoch, data are grouped by saccade type (pro vs. anti); for the visual and saccade epochs, all four task conditions are shown. Data points represent rate changes of individual neurons (positive values denote an increase in activity after cTBS); colors indicate cTBS location (PFC, SHAM, M1). Bars show group medians; black error bars represent SEM. *PFC: prefrontal cortex; M1: primary motor cortex; SEM: standard error of the mean; a.u.: arbitrary units*.

For neurons tuned during the rule epoch (Fig. 7A, D), the linear mixed model revealed no evidence for a main effect of cTBS location, nor any interaction effects including cTBS location on firing rate change (all p > 0.27).

During the visual epoch (Fig. 7B, E), there was no main effect of cTBS location (p = 0.52), but we found a significant interaction between cTBS location and saccade type (F(2,4119) = 18.2, p < 0.001). Specifically, the cTBS-location effects on rate change were due to differences between PFC and M1 (PFC/pro vs. M1/pro, p < 0.001, mean difference = 0.08 ± 0.02; PFC/anti vs. M1/anti, p = 0.005, mean difference = −0.07 ± 0.02) as well as between SHAM and M1 (SHAM/pro vs. M1/pro, p = 0.001, difference = 0.08 ± 0.02; all Bonferroni-corrected). Subsequently, post-hoc analyses of the significant interaction involving cTBS location, saccade type, and monkey revealed that these differences were driven by effects in Monkey Bu (PFC/pro/Bu vs. M1/pro/Bu, p < 0.001, mean difference = 0.17 ± 0.04; M1/pro/Bu vs. SHAM/pro/Bu, p = 0.001, difference = 0.16 ± 0.04).

For the saccade epoch (Fig. 7C,F), the main effect of cTBS location approached significance (F(2,1659) = 2.94, p = 0.053), and post-hoc comparison revealed a small but significant difference between PFC and M1 (t(1659) = −2.42, p_Bonferroni = 0.046; mean difference = −0.02 ± 0.01), while PFC vs. SHAM (p = 0.90) and SHAM vs. M1 (p = 0.74) were not significant. Furthermore, post-hoc analyses of the significant interactions between cTBS location and monkey (F(2,1659) =3.89, p = 0.021) and cTBS location, direction, type, and monkey (F(2,4977)=3.80, p < 0.022) indicated that these effects were largely driven by differences between monkeys Bu and Gr (t(1659)=3.81, p < 0.001, mean difference = 0.03 ± 0.01).

In summary, cTBS appeared to induce only minimal effects in the population of task-modulated activity, with no consistent effect of cTBS location across both monkeys. We also found no evidence for the predicted increase in PFC activity following cTBS-PFC.

#### Session-by-session analysis of cTBS effects on spike rates

Finally, we analyzed the possibility that the neural effects of cTBS may change across days, perhaps being more effective on some days than others. We therefore repeated the above analyses after averaging the changes in spike rate across cTBS across all units recorded on a given day (Figure 8). We observed no main (F(2,42) = 1.20, p = 0.31) or interaction effects (all p > 0.49) of cTBS location on neural activity during the rule epoch. We did observe a significant interaction between cTBS location and saccade type during the visual epoch (Fig. 8B,E; F(2,192) = 7.95, p < 0.001), which was partially driven by a significant difference between SHAM and M1 prosaccades (difference = 0.07 ± 0.02, p = 0.029, Bonferroni-corrected). Additionally, a significant interaction between cTBS location, saccade type, and monkey (F(2,192) = 7.09, p = 0.001) was due in part to differences in prosaccades for Monkey Bu following cTBS over M1 (M1/pro/Bu vs. PFC/pro/Bu, difference = 0.11 ± 0.03, p = 0.045; M1/pro/Bu vs. SHAM/pro/Bu, difference = 0.14 ± 0.03, p = 0.007). Finally, during the saccade epoch, we found a weak but significant interaction for cTBS location x direction x monkey (F(2,272) = 3.19, p = 0.043) where post-hoc results did not survive Bonferroni correction.

**Fig. 8.**
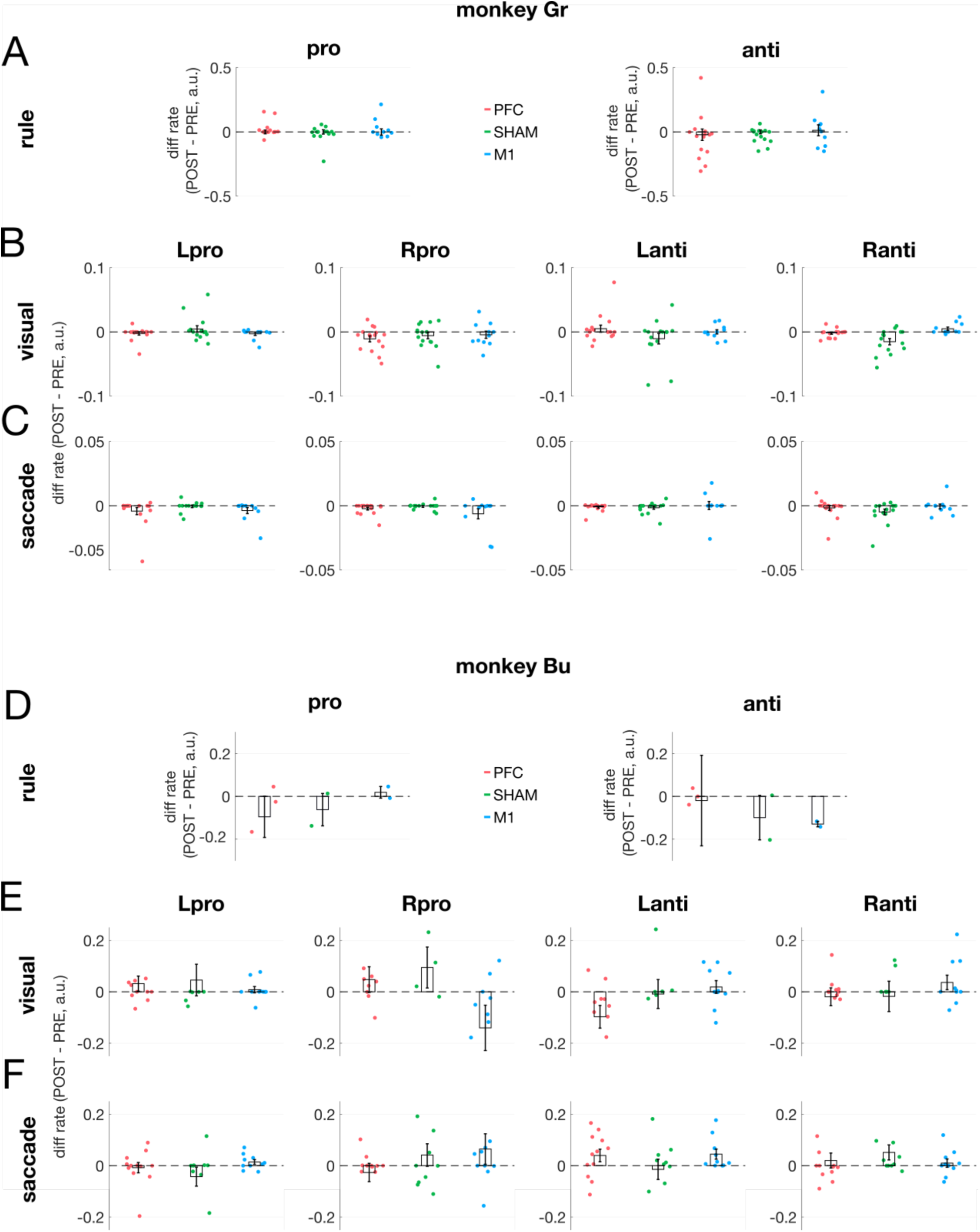
Effects of cTBS on normalized spike rates averaged within sessions. Same format as Fig 8, except that each data point shows the change in spiking activity of significantly task-modulated, averaged within a given session (i.e., one data point per session containing tuned units). *PFC: prefrontal cortex; M1: primary motor cortex; SEM: standard error of the mean; a.u.: arbitrary units*.

Taken together, analyses across both individual neurons and session-level averages revealed no evidence for any consistent influence of cTBS to the PFC on spiking activity in the contralateral PFC. While we observed some significant interaction effects during the visual and saccade epochs, these effects were not consistent across monkeys, often did not survive corrections for multiple comparisons, and varied considerably in both absolute magnitude and directionality. We did not see any evidence for disinhibition following cTBS-PFC when compared to the cTBS-SHAM and cTBS-M1 controls. Broadly speaking, the absence of disinhibitory effects aligns with the lack of any behavioural changes after cTBS.

## Discussion

We investigated the behavioural and neural effects of delivering cTBS to the PFC of two healthy rhesus macaques performing an oculomotor task. Our study tested the hypothesis that cTBS will inhibit the PFC, and disinhibit the mirroring, contralateral PFC. We did not find any reliable behavioral or neural evidence consistent with this hypothesis, and in fact many of our observations went in the direction opposite to these predictions. These are effectively negative results, and we report them in the context of recent reviews that have emphasized the importance of reporting such results for the field (Héroux, Taylor and Gandevia, 2015; Ziemann and Siebner, 2015). Work in animal models, and NHPs in particular, do not afford the opportunity for extensive pilots, and in this Discussion we will consider a number of methodological choices that, in retrospect, may have been suboptimal. Our results contribute to a growing series of studies that question the fundamental assumptions of notion of cTBS-mediated inhibition and consequent disinhibition of callosal targets, and emphasize the challenges associated with assessing changes in neural activity across intermediate timescales given the known fluctuations in PFC activity.

### Choice of behavioural task, and rationale for using a repetitive mode of TMS

We trained our animals to perform an inter-mixed pro- and anti-saccade task that required them to first encode and then remember this instruction in order to correctly complete the trial following peripheral target presentation. The memory component increases the complexity of the task, which is reflected in the elevated pro-saccade error rate particularly in monkey Bu (Fig. 2B) compared to when the instruction cue persists for the entire trial (Everling, Dorris and Munoz, 1998). We chose this task given the excellent understanding of the neural correlates of task performance (Munoz and Everling, 2004), and since anti-saccades have been used in other studies delivering TMS to the FEF in macaques (Valero-Cabre *et al*., 2012) and humans (Olk *et al*., 2006; Beynel *et al*., 2014; Cameron, Riddle and D’Esposito, 2015). Our choice of task is validated by PFC activity reflecting both rule maintenance (Wallis, Anderson and Miller, 2001; Everling and DeSouza, 2005), and the visuo-motor transformation (Bullock *et al*., 2017; Khanna, Scott and Smith, 2020). The interpretation of our results are bolstered through comparisons to results in similar tasks that modulate PFC activity through other means to change behaviour and/or neural activity (Wegener, Johnston and Everling, 2008; Phillips, Johnston and Everling, 2011; Pouget *et al*., 2020).

Another common task used to assess how a given intervention may change neural activity or behaviour is the stimulus-onset asynchrony (SOA) task, which staggers the presentation time of diametrically-opposed targets (Balan *et al*., 2017; Adam, Johnston and Everling, 2019; Kubanek *et al*., 2020). The influence of a given intervention can be assessed by shifts in the timing at which the subject would look to either target with equal likelihood. This is a simpler task than the one we employed, and behavioural shifts in the point of equal selection can often be explained by the changes in RTs to single left or right targets (Rincon-Gonzalez *et al*., 2016). Given the unchanged RTs of either pro- or anti-saccades, it seems unlikely that behavioural effects would have been observed in the SOA task.

A rTMS study necessitates blocked “before-vs-after” analyses. The application of cTBS clearly influenced multiple aspects of oculomotor behaviour and PFC activity, but this influence was largely the same regardless of cTBS target. Simply delivering cTBS likely influences the animal’s r, which temporarily shortens RTs (Fig. 4) and elevates the visual and motor responses (Fig. 7). The dynamics of such changes complicate the identification of effects attributable to cTBS-PFC, as any such changes would have to ride on top of the already substantial changes induced by non-specific arousal. Other groups have reported behavioral and neural changes in NHPs across rTMS that exceed that attributable to non-specific arousal (Romero *et al*., 2019, 2022), but in no cases did we observe results from cTBS-PFC that differed substantially from both cTBS-M1 and cTBS-SHAM. rTMS protocols in humans may be more effective when the targeted area is actively engaged in a task (for a review, see Beynel *et al*., 2019), but the effectiveness and directionality of this approach has not been sufficiently established to warrant the additional training required for experimental animals.

Finally, teaching animals the pro-/anti-saccade rule required progressive stages of training. A nuance in the current dataset is that monkey Bu had a longer training history than monkey Gr before the start of data collection, and in both monkeys the cTBS-M1 sessions were acquired after a substantial number of cTBS-PFC and -SHAM sessions (see Methods, Supplementary Fig 1). This difference in training history likely explains why monkey Gr’s anti-saccade error rates before cTBS were generally lower in the M1 sessions compared to the PFC and SHAM sessions (Fig 2). The importance of training history is also apparent in the work by Gerits and colleagues (2011), wherein saccadic RTs decreased considerably by the time data from cTBS-M1 was collected. But even if we discount the cTBS-M1 data from monkey Gr, the change in error rates across cTBS were remarkably similar for the PFC vs SHAM conditions, and neither exhibited the increase predicted from the hypothesis that cTBS will inhibit neural activity within the targeted area.

### Choice of TMS parameters and controls

#### rTMS protocol

We decided to use cTBS rather than 1-Hz rTMS for both pragmatic and theoretical reasons. Delivery of 600 cTBS pulses requires only ∼40s, whereas an equivalent number of pulses at 1-Hz requires 10 minutes. Given the before-vs-after nature of our analyses, we opted for a briefer time for rTMS delivery. There is also evidence for superior effects of cTBS than 1-Hz rTMS, although such effects may not last as long (Di Lazzaro *et al*., 2010). There is also emerging evidence that delivering more rTMS pulses in a single bout of cTBS results in longer lasting effects (Wischnewski and Schutter, 2015), but doing so would have overheated the TMS coil.

#### Interleaving cTBS sessions

Our decision to not deliver cTBS to PFC or M1 on sequential days was motivated by a desire to avoid possible cumulative effects of cTBS (see Valero-Cabré, Pascual-Leone and Rushmore, 2008). Separating sessions where cTBS is delivered to the same brain area by at least 48 hours permits within-session analysis unconfounded by the history of what was delivered the day before. However, in Monkey Bu we acknowledge that cTBS was delivered to the PFC 24 hours after M1 on a subset of sessions (or vice versa; see Supplementary Fig 1). While we think the degree to which cTBS-M1 influenced PFC activity is fairly minimal (see below), it is possible that this subset of sessions in Monkey Bu featured some degree of a cumulative effect. We avoided cumulative effects in Monkey Gr, and the lack of influence of cTBS across both animals suggests that possible cumulative effects are not having a major impact on our results. This interleaved approach may have decreased the likelihood of seeing cTBS effects, particularly given recent trends advocating for more TMS pulses within a given day, and across sequential days (Lefaucheur *et al*., 2014, 2020). The decision to interleave cTBS sessions is also influenced by the limited number of NHPs, the potential confounds with training history (see above), and the possibility of device failure or decreasing yields over time inherent with chronically-implanted recording arrays. It is also possible that cTBS effects would have been realized had we delivered multiple bouts of cTBS on a given day (e.g., 1200 pulses total; (Gerits *et al*., 2011; Balan *et al*., 2017)), although we note that work in macaques has reported an influence of cTBS on brain activity and behaviour with only 300 individual pulses during 20 s of cTBS (Merken *et al*., 2021; Romero *et al*., 2022).

#### TMS intensity

In humans, a typical pulse intensity for cTBS to M1 is 80% of resting motor threshold (Di Lazzaro *et al*., 2005; Huang *et al*., 2005; for a review see Fitzgerald, Fountain and Daskalakis, 2006). The absence of any effects in our study contrasts with reports of cTBS changing grasping behaviour and neural excitability in the macaque parietal cortex (Merken *et al*., 2021; Romero *et al*., 2022), and on saccade behaviour and intra- and inter-hemispheric functional connectivity in the macaque FEF (Gerits *et al*., 2011; Balan *et al*., 2017). These studies also delivered cTBS at an intensity of ∼80% of resting motor threshold, whereas we used higher pulse intensities above the resting motor threshold to resemble that used in the DLPFC for treatment of intractable depression (Lefaucheur *et al*., 2014; Van Rooij *et al*., 2024). Dose–response relationships in TMS are often nonlinear (Bergmann and Hartwigsen, 2020), and the tradeoff between pulse intensity and effect focality is particularly pertinent in smaller brains, but we view it as unlikely that a higher pulse intensity explains why the observed behavioral or neural results of cTBS did not conform to predictions.

We are also confident that cTBS of the PFC or M1 influenced brain activity. We did not record neural activity in the PFC targeted by cTBS, but we confirmed on cTBS-PFC sessions that the reliable and distinct contralateral thumb twitches evoked by single pulse TMS to M1 were less pronounced and less frequent for single-pulse TMS to PFC, consistent with a high degree of focality. Many aspects of our implant design and pulse intensity resemble that used in our previous study, wherein single pulses of TMS over a widespread area of frontal cortex, including the PFC location we targeted, induced a feed-forward neck muscle response (Gu and Corneil, 2014). In retrospect, obtaining such objective markers to single-pulse TMS-PFC would have provided further insights about the effects of cTBS. For example, Romero and colleagues (2022) showed that cTBS to the parietal cortex reduced the neural response to single-pulse TMS. Despite such a reduction, cTBS to the parietal cortex did not influence the feedforward response to a visual stimulus, paralleling our finding of the lack of any influence of cTBS-PFC on the visual response in the contralateral hemisphere.

#### Choice of controls

We included both a SHAM and a brain (M1) control to control for non-specific effects of cTBS delivery, including tactile and auditory sensations, and changes in arousal. This SHAM control is not ideal, as we did not have access to a true sham coil. Instead, our SHAM control relies on the additional physical separation induced by moving the TMS coil a further 2 cm above the acrylic, which itself lies ∼5 mm above the intact skull. While this 2 cm distance may seem modest relative to the electric field estimates of standard figure-8 coils used in human research (Roth *et al*., 2007; Deng, Lisanby and Peterchev, 2013), the 25 mm figure-8 coil used here has a higher degree of focality but consequently smaller depth of penetration (Deng, Lisanby and Peterchev, 2013). Thus, while we cannot exclude the possibility that our SHAM condition completely eliminates the possibility of influencing PFC activity, the degree of activation will certainly be less than that achieved by cTBS-PFC.

Considerations of focality also pertain to the suitability of our brain control, where we delivered cTBS to M1. While the PFC and M1 cTBS locations were separated by ∼8mm, distinct neck muscle responses can be evoked from these locations (Gu and Corneil, 2014). However, given that our stimulation intensity is above resting motor threshold, is it possible that both cTBS-PFC and cTBS-M1 influenced the oculomotor network? While we cannot exclude this possibility, our results provide no consistent evidence for this across both NHP, as we observed no evidence for any ‘dosing’ like effects, wherein the behavioral or neural effects of cTBS-M1 could have been greater than cTBS-SHAM, given closer proximity of the former to the PFC.

Finally, an alternative approach may have been to use an intensity control (e.g., delivering cTBS-PFC at ∼80% rather than ∼110% of motor threshold) to investigate potential dose-dependent effects. While a reasonable approach, the controls we did use may actually serve as more extreme versions of this type of control, if indeed such controls induced unintended influences on the PFC. Regardless, all of these possibilities must be interpreted in light of the absence of clear differences between cTBS applied to the PFC, M1, and SHAM conditions, and the lack of evidence for any behavioral or neural effects that conformed to the predictions arising from our hypotheses.

#### Assessment window for behavioural and neural effects

Is it possible that our analyses examined intervals that were either too soon or too late after cTBS? The original report showed that the decrease in cortical excitability following cTBS peaks ∼5-10 minutes later (Huang *et al*., 2005), but decreases certainly begin within the first few minutes. More recent reviews have focused on the duration of such effects on neural excitability, which exceed 20 minutes or more depending on the number of cTBS pulses delivered (Thut and Pascual-Leone, 2010; Wischnewski and Schutter, 2015). In the macaque, the work from the Janssen lab (after 300 pulses of cTBS) show more variability in the timing of peak effects after cTBS to the parietal cortex, with such effects peaking ∼20-30 min after cTBS (Merken *et al*., 2021; Romero *et al*., 2022). However, even in this work, reliable changes in behaviour or neural excitability begin to appear soon after cTBS, even if the peak effect is delayed. Our presentation of saccadic RT (Fig. 4) or neural activity (Fig. 6) through time emphasizes that the time-dependent effects were common to all cTBS conditions. Finally, we note that mechanisms often invoked to explain the delayed nature of the effects of cTBS, such as a delayed increase in GABA levels (Stagg *et al*., 2009) in the stimulated area and an associated decrease in the contralesional hemisphere (Matsuta *et al*., 2022) should have altered saccade behaviour in predictable ways, given the effects of delivery of GABA agonists or antagonists to the FEF (Dias, Kiesau and Segraves, 1995). For example, increased GABA in the stimulated (rightward) hemisphere should have increased the RT and decreased the peak velocity of leftward saccades, whereas decreased GABA in the opposite (leftward) hemisphere should have decreased the RT and increased the peak velocity of rightward saccades. We found no evidence of this in our cTBS-PFC data.

#### Further technical considerations

The TMS coil used for this study has been widely employed in previous NHP research (e.g., Amaya *et al*., 2010; Gerits *et al*., 2011; Valero-Cabre *et al*., 2012; Gu and Corneil, 2014; Romero *et al*., 2019, 2022; Merken *et al*., 2021). It is big enough for efficient delivery of cTBS while small enough given the spatial constraints imposed by cranial implants. The cranial implant was optimized to avoid field distortion and minimize the distance from coil to skull by incorporating a thin acrylic layer (∼5 mm; for details, see (Gu and Corneil, 2014)). Precise and repeatable coil placement was ensured via neuronavigation, which was further confirmed via the consistency of the evoked thumb twitches from single-pulse TMS-M1 at the start of every session; coil orientation has been shown to be a relatively negligible factor for this type of coil (Alekseichuk *et al*., 2019). Overall, it is unlikely that any of these technical considerations explained the lack of cTBS-PFC effects. Finally, we delivered biphasic pulses, while some evidence is showing that monophasic pulses induce stronger and longer-lasting effects (Tings *et al*., 2005; De Lima-Pardini *et al*., 2023); this may be a parameter to modify in future studies.

### No selective influence of cTBS-PFC on spiking activity in the mirroring, contralateral PFC

Neural activity was not recorded from the stimulated area, due to concerns that TMS would negatively impact the stability or functioning of the chronically-implanted Utah array. However, the effects of non-invasive brain stimulation are not limited to the targeted area, and the possibility that cTBS-mediated inhibition can be used to induce balanced or compensatory excitation in the mirroring, contralateral target has clear value for basic and clinical research. One message of our manuscript is the lack of evidence for this in terms of task-relevant spiking activity in the contralateral PFC in healthy macaques. This absence is all the more surprising considering findings in macaques that show that very similar cTBS protocols can alter the excitability of the targeted area to single-pulse TMS in the parietal cortex (Romero *et al*., 2019), and functional connectivity in the frontal cortex, including to the mirroring, contralateral target (Balan *et al*., 2017). In our study, cTBS-PFC may not have induced a strong enough perturbation for changes in task-related neural activity to stand out from the already substantial fluctuations in neural activity in this area. Further, there may well be other perhaps more sensitive measures of neural activity (e.g., local field potentials, and/or spike-field coherence) that do change selectively with cTBS-PFC; such measures will be explored in a future manuscript.

PFC may also not be an ideal location to test the hypothesis of cross-hemisphere disinhibition (Pascual-Leone, Walsh and Rothwell, 2000). There is some evidence that the oculomotor network is fairly resilient to perturbations, perhaps because animals execute visually-guided saccades during the inter-trial interval; thus, it may only be in the case of more potent or widespread lesions that behavioural deficits emerge, and even then any deficits can still be surprisingly task-dependent (Peel *et al*., 2017, 2020; Vaidya *et al*., 2019). It is also possible that effects would have been seen had cTBS been combined with a pre-existing lesion, and our failure to observe cTBS-PFC effects in healthy animals may not generalize to animal models of clinical conditions, such as stroke. Finally, a recent meta-review (Kirkovski *et al*., 2023) on cTBS protocols emphasized that cTBS-PFC effects are more variable than cTBS-M1, perhaps due to inherent variability in both task-related activity and in PFC activity more generally (Cowley *et al*., 2020; Khanna, Scott and Smith, 2020). Such variability in PFC activity may thus be a doubled-edge sword that influences both the ability of cTBS to influence outcome measures, and the ability to differentiate such outcome measures from noise.

Finally, and inherent to many NHP studies, our study’s sample size of only two NHPs limits the generalizability of the results and may not capture individual variability in cTBS response. It is possible that the two NHPs in this study were simply non-responders. While non-responder rates in NHPs have not been documented to our knowledge, non-responder rates in humans to TBS range between 30% to 50% (López-Alonso *et al*., 2014; Huang *et al*., 2017; Boucher *et al*., 2021; Ozdemir *et al*., 2021). If these rates generalize to NHPs, then the chances that both our NHPs are non-responders is ∼10% to 25%. However, evidence in humans from M1 suggests that a given individual may respond to TBS on some days but not others (Perellón-Alfonso *et al*., 2018; Boucher *et al*., 2021). Therefore, the limitations of our small sample size is counterbalanced by repeated sampling of the same subject across many days. Finally, analyses motivated by the possibility of the effectiveness of cTBS on some days but not others failed to reveal anything unique for cTBS-PFC (Fig 4, Table 1).

### Conclusions

Our findings offer no support for the widely held assumptions that cTBS, at least of the PFC in a healthy brain, will inhibit neural activity within the stimulated area, and consequently disinhibit neural activity in the mirroring, contralateral target. Our results challenge the simple ‘rebalancing’ logic often applied for cTBS in the healthy brain, and are in line with recent work showing limited reliability and reproducibility after cTBS of M1. More work is needed to better understand the effects of repetitive modes of TMS, including cTBS, on brain and behaviour in human and animal models.

## Supplementary Materials

**Fig. S1.**
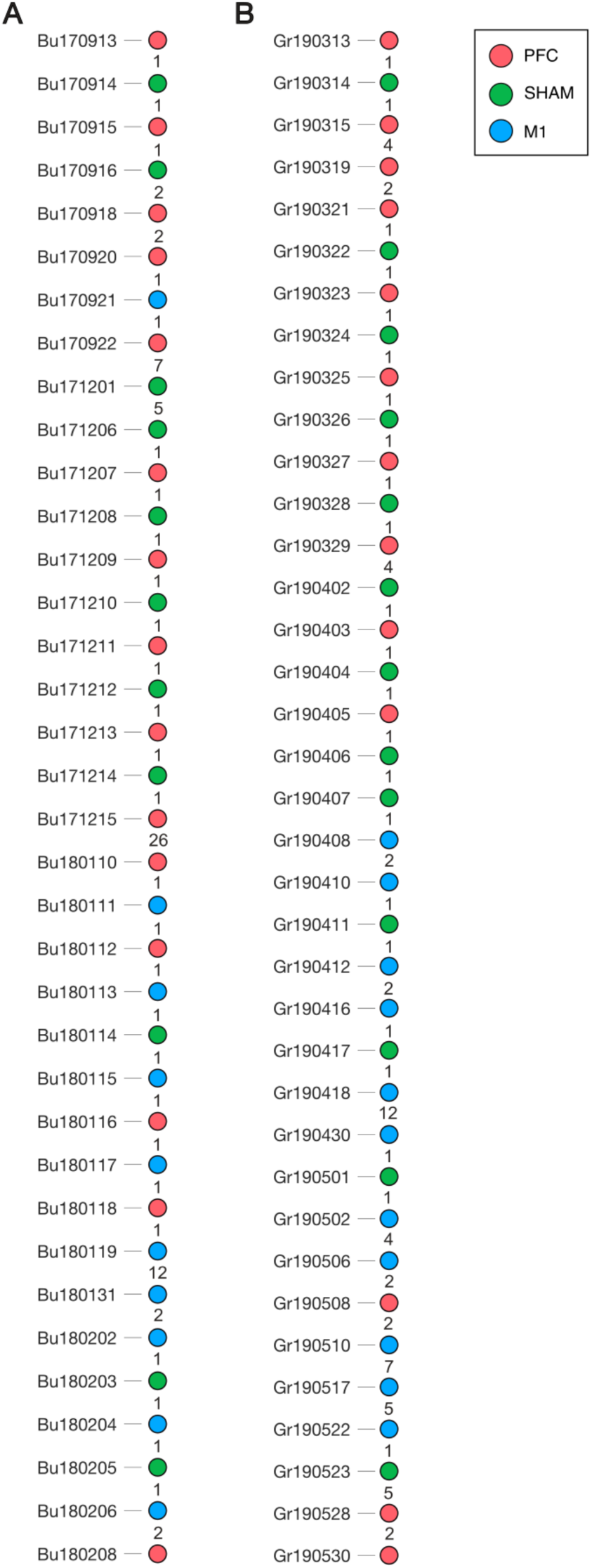
Timeline of Recording Sessions and Stimulation Site Mapping for Monkeys Bu (A) and Gr (B). Dot colors indicate cTBS target location; numbers between dots reflect the number of days between sessions. Dates follow the format: [MonkeyIDYYMMDD]. *PFC: prefrontal cortex; M1: primary motor cortex*.

**Fig. S2.**
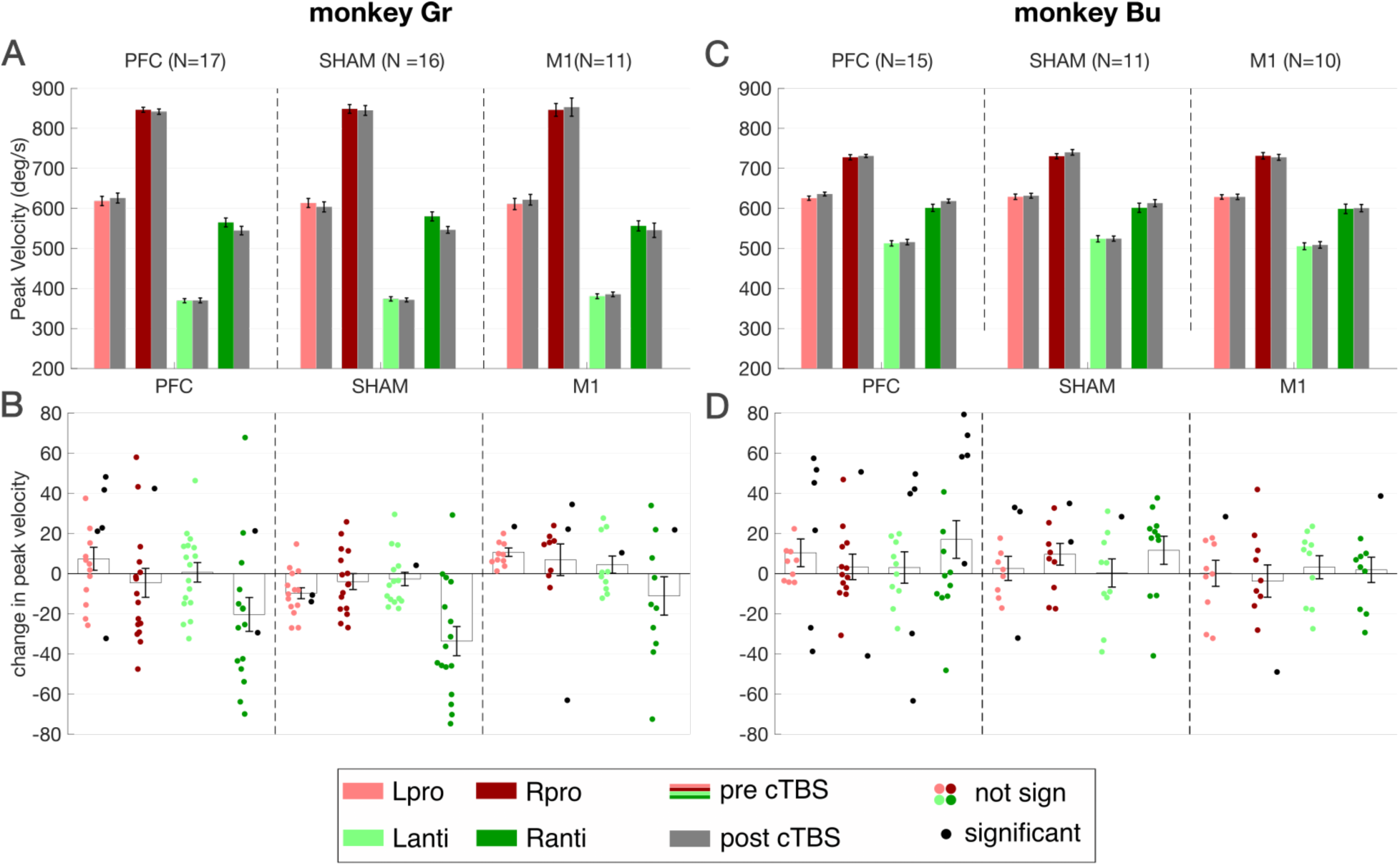
Effects of cTBS on saccadic peak velocity. Same general format as Figs. 2 and 3. **A,B**. Peak velocities before and after cTBS-delivery or both monkeys, averaged across sessions. **C,D**. Change in peak velocity following cTBS delivery, relative to baseline. Black or coloured markers shifted to the right or left denote individual sessions with significant on non-significant changes in peak velocity, respectively (1-way ANOVA, uncorrected). *PFC: prefrontal cortex; M1: primary motor cortex;*

### Task-related effects identified by Linear Mixed Model Analyses

This section lists the results from our Linear Mixed Model (LMM) analyses for task-related effects, excluding the factor cTBS location.

#### Error Rate changes (Figure 2)

Our analysis of error rates revealed a significant main effect of Saccade Type (F(1,222)=8.3, p=0.004) and a significant interaction between Saccade Type and Monkey (F(1,222)=8.7, p=0.004).

#### Reaction time differences (Figure 3)

For saccadic reaction times (RTs), we found significant main effects of Saccade Type (F(1,222)=42.6,p<0.001) and Monkey (F(1,74)=17.9,p<0.001). We also observed several significant interactions for Saccade Direction and Monkey (F(1,222)=6.2, p=0.013), Saccade Type and Monkey (F(1,222)=20.5, p<0.001), as well as a significant three-way interaction among Saccade Direction, Saccade Type, and Monkey: F(1,222)=6.2, p=0.013.

#### Short-term reaction time differences (+/− 25 trial, Figure 4)

The LMM on RTs within the +/− 25 trial window identified significant main effects for Block (F(1,15921)=558.2, p<0.001), Saccade Type (F(1,15921)=3397.6, p<0.001), Saccade Direction (F(1,15921)=195.2, p<0.001), and Monkey (F(1,495)=377.7, p<0.001).

Numerous significant interactions were also found:

- Monkey x Block: F(1,15921)=264.8, p<0.001
- Monkey x Direction: F(1,15921)=20.0, p<0.001
- Monkey x Type: F(1,15921)=1507.6, p<0.001
- Monkey x Block x Direction: F(1,15921)=16.6, p<0.001
- Block x Direction x Type: F(1,15921)=3.8, p<0.05
- Block x Direction x Type x Monkey: F(1,15921)=24.3, p<0.001

#### Peak Velocities (Figure S1)

For peak saccade velocities, we observed significant main effects of Saccade Direction (F(1,222)=4.9, p=0.029), Saccade Type (F(1,222)=4.4, p=0.037), and Monkey (F(1,74)=5.6, p=0.021). Significant two-way interactions were found between Monkey and Saccade Direction (F(1,222)=14.1, p<0.001) and Monkey and Saccade Type (F(1,222)=10.4, p=0.001), as well as a significant three-way interaction among Monkey, Saccade Direction, and Saccade Type (F(1,222)=10.8, p=0.001).

### Spike Rate Difference, Tuned Units (Figure 7)

- **Cue epoch:** The analysis of firing rates for tuned units revealed no significant effects during the cue epoch.
- **Visual epoch:** During the visual epoch, we found a significant interaction between Saccade Direction and Saccade Type (F(1,4119)=6.7, p=0.01).
- **Saccade epoch:** For the saccade epoch, there was a significant main effect of Monkey (F(1,1659)=14.5, p<0.001).

### Spike Rate Difference, Sessions (Figure 8)

- **Cue epoch:** In the session-level analysis, we found a significant main effect of Monkey during the cue epoch (F(1,42)=5.61, p=0.023).
- **Visual epoch:** No significant effects during the visual epoch.
- **Saccade epoch:** During the saccade epoch, a significant main effect of Monkey

**Supplementary Video S1.**
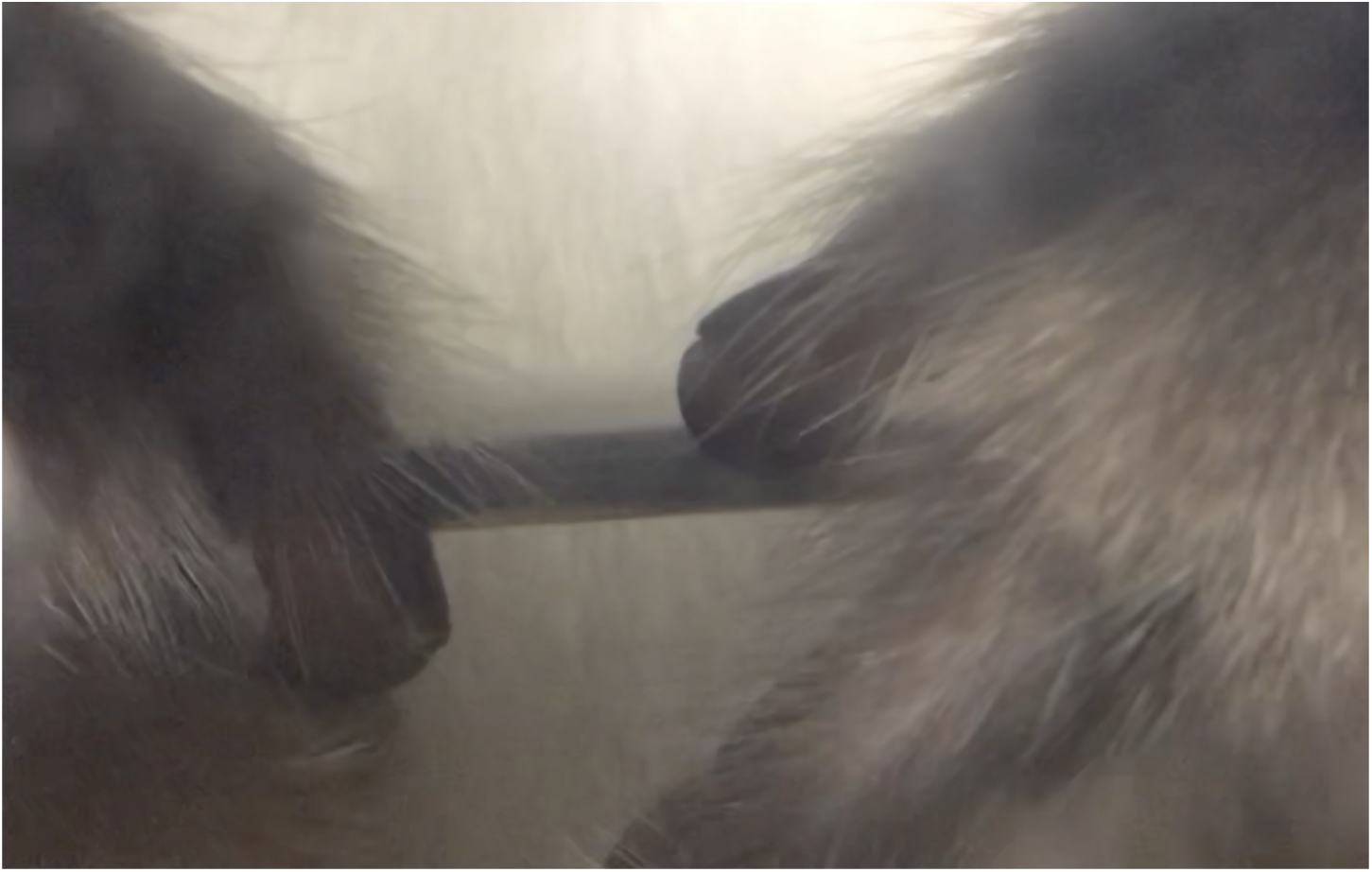
Single-pulse TMS over the hand area of right M1 in awake monkey Bu reliably evoked highly localized contra-lateral thumb twitches. The example shown in this video confirms the suitability of the neuronavigation approach prior to cTBS (see methods). The single pulses in this video (audible as distinct clicks) were delivered at 30% of stimulator output, matching the intensity used for cTBS over the same site (M1) or PFC in subsequent recording sessions. Note the specificity of the motor response to the thumb, irrespective of hand and arm posture. *PFC: prefrontal cortex; M1: primary motor cortex;*

## Acknowledgements

This work was supported by Operating Grants from the Canadian Institutes of Health Research (CIHR; MOP-123247, MOP-142317, PJT-186160). We thank Dr. Julio Martinez-Trujillo for surgical expertise during the insertion of the Utah arrays.

